# Dynamic mode decomposition for analysis and prediction of metabolic oscillations from time-lapse imaging of cellular autofluorescence

**DOI:** 10.1101/2025.03.06.641958

**Authors:** Daniel Wüstner, Henrik Helge Gundestrup, Katja Thaysen

## Abstract

Metabolic oscillations are a common phenomenon in cell biology. They are based on non-linear coupling of biochemical reactions and can show rich dynamic behavior including sustained and damped oscillations, as found, for example, in glycolysis of yeast and other eukaryotic cells. Metabolic oscillations are often studied by time-lapse imaging of cellular autofluorescence based on the changing abundance of NAD(P)H, but the analysis of such experimental data is challenging. Here, we show that dynamic mode decomposition (DMD), a numerical algorithm for linear approximation and spectral analysis of non-linear dynamics, allows for dissecting glycolytic oscillations in simulations and experiments in a fully data-driven manner. By combining DMD with time-delay embedding the spatiotemporal dynamics of sustained and damped glycolytic oscillations can be learned. Together with a rigorous assessment of spurious eigenvalues, via residual DMD, this provides a unique spectrum for each scenario, allowing for high-fidelity time-series and image reconstruction as well as for phenotyping different starvation conditions. The ability of DMD to predict future time points depends on the delay embedding dimension and is comparable to that of long short-term memory (LSTM) neural networks. Together, our results demonstrate the potential of DMD for analysis of time-lapse microscopy of metabolic oscillations in living cells.

## Introduction

The energy metabolism of eukaryotic cells is tightly controlled to meet the demands of a constantly changing environment. Oscillations in abundance of metabolites are a hallmark of cellular respiration in various cell types [1, 2]. They have been studied in great detail in baker’s yeast, *Saccharomyces cerevisiae*, in which oscillations occur due to fluctuations in concentrations of key metabolites accompanied by oscillations in mitochondrial membrane potential [3–5]. Glycolytic oscillations can be monitored by label-free measurements of the intrinsic fluorescence of the reduced form of the electron carriers nicotinamide adenine dinucleotide (phosphate) (NAD(P)H) with an excitation around 360 nm and emission between 400-500 nm [5, 6]. Alternatively, NAD(P)H can be excited at 720 nm by a two-photon process, which is particularly suitable for mammalian cells and tissues, whose thickness prevents high-contrast imaging by one-photon excitation [7–9]. Once oxidized, the fluorescence signal vanishes, while reformation of the reduced form, NAD(P)H, gives again a fluorescent signal, such that oscillations in autofluorescence around 450 nm monitor the interconversion of this electron carrier between the reduced and oxidized state.

While the nicotinamide adenine dinucleotide phosphate (NADPH) is the main electron carrier in anabolic reactions, nicotinamide adenine dinucleotide (NADH) serves the same role in catabolic processes, such as glycolysis. Baker’s yeast uses glucose as primary energy source during exponential growth, and a large body of work has been devoted to the analysis of glycolytic oscillations by monitoring NADH fluorescence over time [10]. In synchronized yeast populations, time-varying NADH concentrations of individual cells oscillate in-phase, allowing one to measure glycolytic oscillations spectrophotometrically [4, 5]. In contrast, imaging-based analysis allows for monitoring individual cells, thereby revealing partial coherence phenomena as well as desynchronization dynamics and even glycolytic waves traveling throughout the imaged cell population [11–13]. Various computational methods have been applied to the analysis of such time-lapse video data [14, 15], but such tools are often designed for a particular condition only [11, 13, 16].

Here, we present the application of dynamic mode decomposition (DMD) to the analysis of glycolytic oscillations in yeast cells based on time-lapse imaging of cellular autofluorescence of NADH fluctuations. DMD is a projection-based computational technique, which provides a linear approximation of non-linear dynamics in a purely data-driven manner [17]. DMD aims to describe complex spatiotemporal signals by a finite-dimensional approximation of the Koopman operator using only measured snapshots of the dynamics as input. This allows for approximating non-linear dynamics by a linear matrix decomposition, providing dynamic modes separated by the individual time scales of the underlying dynamics. DMD has been widely applied to imaging data, including separation of background from foreground, object tracking, video shot detection, motion correction, image segmentation, disease diagnostics in biomedical imaging and detection of intracellular protein aggregates from live-cell microscopy data [18–24]. The performance of DMD can be greatly improved by making use of delay embedding, which means that DMD is applied to time-shifted versions of the original data, thereby enriching the data with past information for improving the reconstruction capabilities [17, 25–27]. This method is based on Taken’s embedding theorem, which states that for a given time series sampled from a high-dimensional attractor, one can approximate the attractor using time-shifted versions of the original time series [27, 28]. This approach has been implemented in several variants of DMD including higher order DMD (HoDMD) and Hankel alternative view of Koopman (HAVOK) analysis, for example in the context of analysis of fluid dynamics but also for reconstructing 3D and polarization dependent fluorescence microscopy data [25, 26, 29, 30]. We show here that time-delay embedding allows for accurate reconstruction of simulated and experimental glycolytic oscillations, and we assess the extent of time delay needed for achieving faithful results. We also show that DMD’s ability to predict future time points depends on the delay-embedding dimension and is on par with that of Long-Short-Term-Memory (LSTM) networks. Together, our results provide a guide in using DMD for analysis of biological time series data gathered by time-lapse imaging of living cells.

## Results

### Metabolic oscillations exemplified by feedback control of glycolysis are inferred by HoDMD

Kinetic models of glycolysis in eukaryotic cells are primarily based on ordinary differential equation (ODE) systems, which can become rather complex depending on the purpose of the analysis [31–34]. For the sake of our analysis, we restrict the simulations of glycolysis to a minimal model, able to model glycolytic oscillations. Our model is a version of the one developed by Higgins (1964) and is based on a positive feedback due to allosteric activation of phosphofructokinase (PFK) by fructose-1,6-bisphosphate (F16BP) [3, 35]. This can be modeled by a simple autocatalytic reaction network of the form shown in Fig. 1A, leading to the following system of non-linear ordinary differential equations:

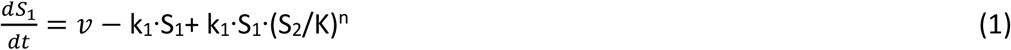

**Figure 1.**
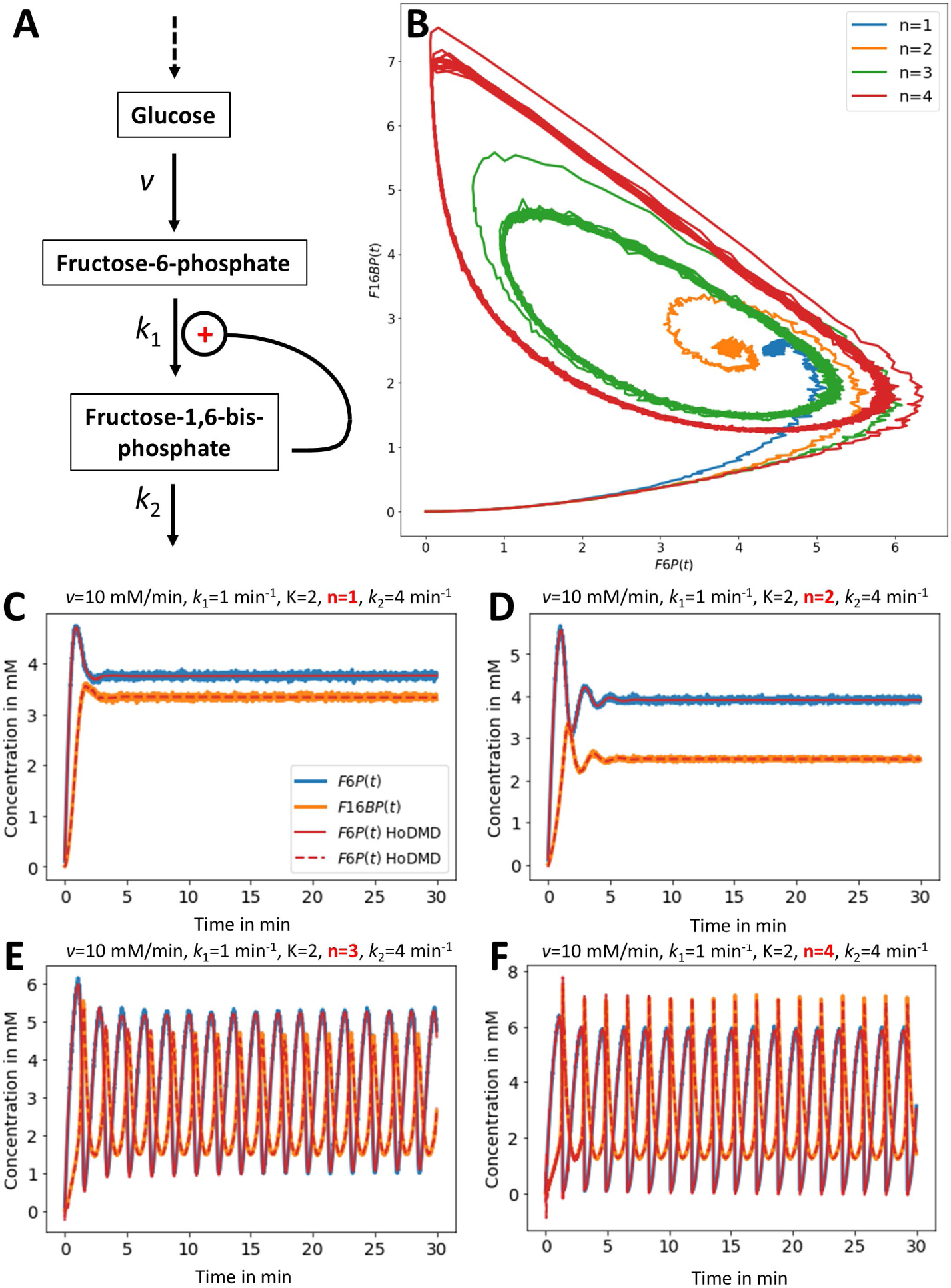
Minimal model of glycolysis and its analysis using higher-order DMD. A steady state model of glycolysis has been formulated based on positive allosteric regulation of phosphofructokinase by its product fructose-1-6-bisphosphate (F16BP). In this model, glucose is supplied with constant influx v, while the concentrations of fructose-6-phosphate (F6P) and of F16BP are allowed to vary (A). A positive feedback indicated by ‘+’ stimulates the conversion of F6P to F16BP followed by constant reaction flux, removing F16BP during the following steps of glycolysis. This model produces different phase portraits depending on the extent of cooperativity of the feedback loop, as shown in panel (B); damped oscillations for n=1 and 2 but a limit cycle with sustained oscillations for high cooperativity, i.e., n=3 and 4. (C-F) simulated time courses for F6P (blue) and F16BP (orange) and their reconstruction by HoDMD in straight and dashed red lines, respectively, are shown for n=1 (C), n=2, (D), n=3 (E) and n=4 (F). All other parameters are as indicated on top of panel C-F.

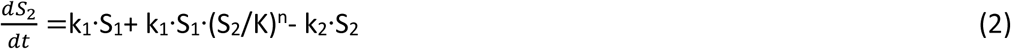

Here, S_1_ and S_2_ are the concentrations of fructose-6 phosphate (F6P) and F16BP, respectively. The reaction rate for the first step, v, is constant and describes the constant inflow of glucose into the system. Thus, this step resembles both the import of glucose and its conversion into glucose-6-phosphate by hexokinase in the presence of constant ATP concentration [36]. While this is a strong simplification of real yeast glycolysis, it allows us to simulate the key characteristics of glycolysis with a minimal number of parameters. The model simulations are intended to provide synthetic data for assessing the performance of DMD in describing the resulting complex dynamic behavior. We do not claim, that the mechanisms used here are the ones which actually cause glycolytic oscillations, in particular in yeast cells, which later studies showed to be the consequence of varying concentrations of various metabolites including NADH/NAD^+^, the ATP/AMP ratio and of acetaldehyde [31, 32, 34]. In fact, F16BP is more important in allosterically activating pyruvate kinase in yeast glycolysis, thereby linking formation of the end product of glycolysis to fermentation and oxidative phosphorylation.

In our toy model of allosteric activation of PFK, the conversion of F6P into F16BP is described by k_1_, the rate constant for the PFK reaction, *n* is the Hill coefficient describing the extent of cooperativity of allosteric activation of PFK by F6P and K is the equilibrium constant for this binding step. The second reaction is modeled as linearized Michaelis-Menten reaction, in which k_2_ is the rate constant for the aldolase reaction. This system can give rise to damped oscillations for n=2 and sustained oscillations for n ≥ 3 and for *k*_2_ =4 min^-1^ (Fig. 1B-F). This is also illustrated in the phase portrait of the two reactions in Figure 1B, for which the system of Eqs. 1A and B were simulated with some additional noise to emulate experimental conditions (Fig. 1B, see Materials and methods for details).

In this case, the positive feedback due to allosteric activation of PFK by F16BP will cause a continuous increase in production of S_2_ (i.e., F16BP), until S_1_ (i.e., F6P) is depleted. This causes a decline in the concentration of F16BP until more F6P due to conversion from G6P is available, and the cycle can begin again. If the aldolase step is slower, i.e. *k*_2_ =3 min^-1^, only damped oscillations are found, independent of the cooperativity of the feedback step, since this step separates regions of stable and unstable steady state (see below and [36]). The initial conditions for these simulations are S_1_(t=0)=S_2_(t=0)=0, so the obtained oscillations solely occur due to the constant inflow and outflow from the system. One also sees, that the oscillations of F16BP are delayed (phase-shifted) compared to those of F6P, which reflect the sequential reaction scheme of glycolysis. This has been observed before, as reviewed and discussed in Heinrich and Schuster (1996) [36]. For stronger non-linearity of the positive feedback (n≥3) and intermediate levels of glucose influx, the steady state of the system shows sustained oscillations and a limit cycle in the phase portrait with a fixed phase shift between both metabolites (Fig. 1B, E and F).

In order to analyze this data by DMD, we apply time-delay embedding, a method which has been shown to approximate the true Koopman eigenfunctions and eigenvalues in case of sufficient snapshot data [37]. The key idea of delay embedding is to increase the available data set by calculating time-shifted versions of the original data resembling past observations [28, 38]. This concept can be illustrated by plotting the time course of F6P for a given parameter set against its time-shifted version, where one finds, that the attractor of Fig. 1B can be approximated depending on the chosen time-shift *d* (Fig. S1). The approximated attractor, sometimes also called a ‘shadow manifold’, M’, is diffeomorphic to the original manifold, M, but only for certain values of the delay, *d* [38–40]. The conditions for obtaining a faithful reconstruction of the original attractor have been studied in detail, and it was found that delay-embedding requires a one-to-one map of the original manifold, M, to the shadow manifold M’, with no two points mapped to the same point [40]. As shown in Fig. S1, if the delays are too closely spaced relative to the periodicity of the oscillations, different points of the system’s cycle can map to the same point, causing overlap and failure to preserve the system’s topological structure.

Time-delay embedding has been extensively used for state-space reconstruction, since it allows for characterizing the dynamic system, even if only partial measurements of the state-space are available [39, 40]. Upon creating time-shifted versions of the observed dynamics, the augmented data set can be analyzed using DMD, thereby allowing for finding better eigenfunctions to approximate the invariant subspace of the Koopman operator compared to standard DMD [25, 26, 29, 37]. There exist several approaches to include past information in the DMD analysis. One popular approach is to produce a snapshot matrix augmented with time-shifted versions of the time series data as consecutive rows, a so-called Hankel matrix, onto which standard DMD is applied [29, 37, 41]. Another approach is to approximate higher-order derivatives of the data using shifted versions of the original data [25]. First, we applied the latter method, called higher-order dynamic mode decomposition (HoDMD), as implemented in the PyDMD python package, to the simulated glycolytic oscillations [42]. We find that HoDMD can faithfully reconstruct this complex dynamic behavior for varying degrees of non-linearity (Fig. 1C-F, red lines). More specifically, we find that the reconstructed time courses obtained with a delay constant of *d*=300 exactly coincide with the simulated ones, both, in case of damped and sustained oscillations. The computed eigenvalues are shown on the unit circle for the first species (i.e., F6P) revealing an increasing contribution of imaginary values, the more oscillatory the system becomes for higher values of the feedback cooperativity, n (Fig. S2A-D). A very similar behavior was found for the eigenvalues of the second species (i.e., F16BP) with HoDMD, while standard DMD failed to dissect the dynamics of the system properly (not shown). We conclude that time-delay embedding is instrumental for proper decomposition of sustained and damped oscillations in a non-linear biochemical reaction system by DMD. Furthermore, we show that the HoDMD reconstruction efficiently smoothens the time traces and is more accurate than Fourier filtering, particularly for damped oscillations (Fig. S2E-F). The good performance of DMD with delay embedding in denoising and reconstructing data confirms our earlier studies using time-lapse and 3D microscopy data [24, 43].

### Stability analysis of the two-state glycolysis model

An advantage of the simple two-state glycolysis model is that its system behavior can be analyzed analytically. By setting Eqs. 1 and 2 to zero, one finds expressions for the steady state concentration of F6P and F16BP (see Appendix, Eqs. A1 and A2). The stability of the dynamic system can be assessed by a Taylor expansion around this steady state and truncating the series after the first term leading to the Jacobian matrix of the system [36, 44]. The Jacobian of this simplified glucose model was solved analytically and is given in Eqs. A3 to A7 of the Appendix. The eigenvalues of this matrix are derived in Eqs. A8 and A9. They are plotted for different values of *k*_2_, the rate constant for the aldolase reaction, as a function of glucose influx for a Hill coefficient of n=3 in Fig. 2A-D. For *k*_2_ =3 min^-1^, and either low (*v*_0_ < 7.5 mM/min) or high ((*v*_0_ > 12.5 mM/min) glucose influx the eigenvalues have both negative real parts, showing that the steady state is stable for these cases. Thus, after a small perturbation, the dynamics of the system will asymptotically return to the steady state for these parameter values. In contrast, for intermediate glucose concentrations, the system is unstable, as inferred from the eigenvalues having positive parts (Fig. 2A and C). In general, a steady state of such a two-component system is stable as long as the trace of the Jacobian is negative, while its determinant is positive [36, 44]. One can see from Fig. 2E-G and Eq. A10, that the determinant of the Jacobian is positive for all parameter combinations, while its trace changes sign for intermediate glucose influx, which is more pronounced for *k*_2_=4 min^-1^ compared to *k*_2_=3 min^-1^. This change of sign of the trace of the Jacobian is characteristic for a Hopf bifurcation, which happens at the bifurcation pint of n=2.4 for this system [36, 44]. For positive trace of the Jacobian, which happens for intermediate glucose influx, high cooperativity of the feedback and fast aldolase reaction (i.e., *k*_2_=4 min^-1^), the steady state becomes unstable, and from the plot of trace squared minus four times the determinant, one sees that an unstable spiral forms under those conditions (Fig. 2E-G). This is the region, in which the system establishes a limit cycle, i.e. sustained oscillations in the abundance of both metabolites. In contrast, for *k*_2_=3 min^-1^ and glucose ≥ 10 mM/min, a stable focus (also called a stable spiral) forms with damped oscillations (Fig. 2D-F). For high glucose influx (i.e., *v* ≥ 17 mM/min), the stable focus is found for fast aldolase reaction (i.e., *k*_2_=4 min^-1^), but the dynamic behavior changes to a stable node for slower aldolase reaction (i.e., *k*_2_=3 min^-1^, Fig. 2D-F). The latter is characterized by the absence of oscillations. The limit cycle observed for n=3 and n=4 is in accordance with the Poincare-Bendixson theorem, which states, that as long as the trajectories of the species are bounded on the manifold M for t → ∞, a closed cycle is formed [36, 44]. The corresponding eigenvalues reflect the change in system behavior for increasing feedback cooperativity and glucose influx (Fig. 2A-D).

**Figure 2.**
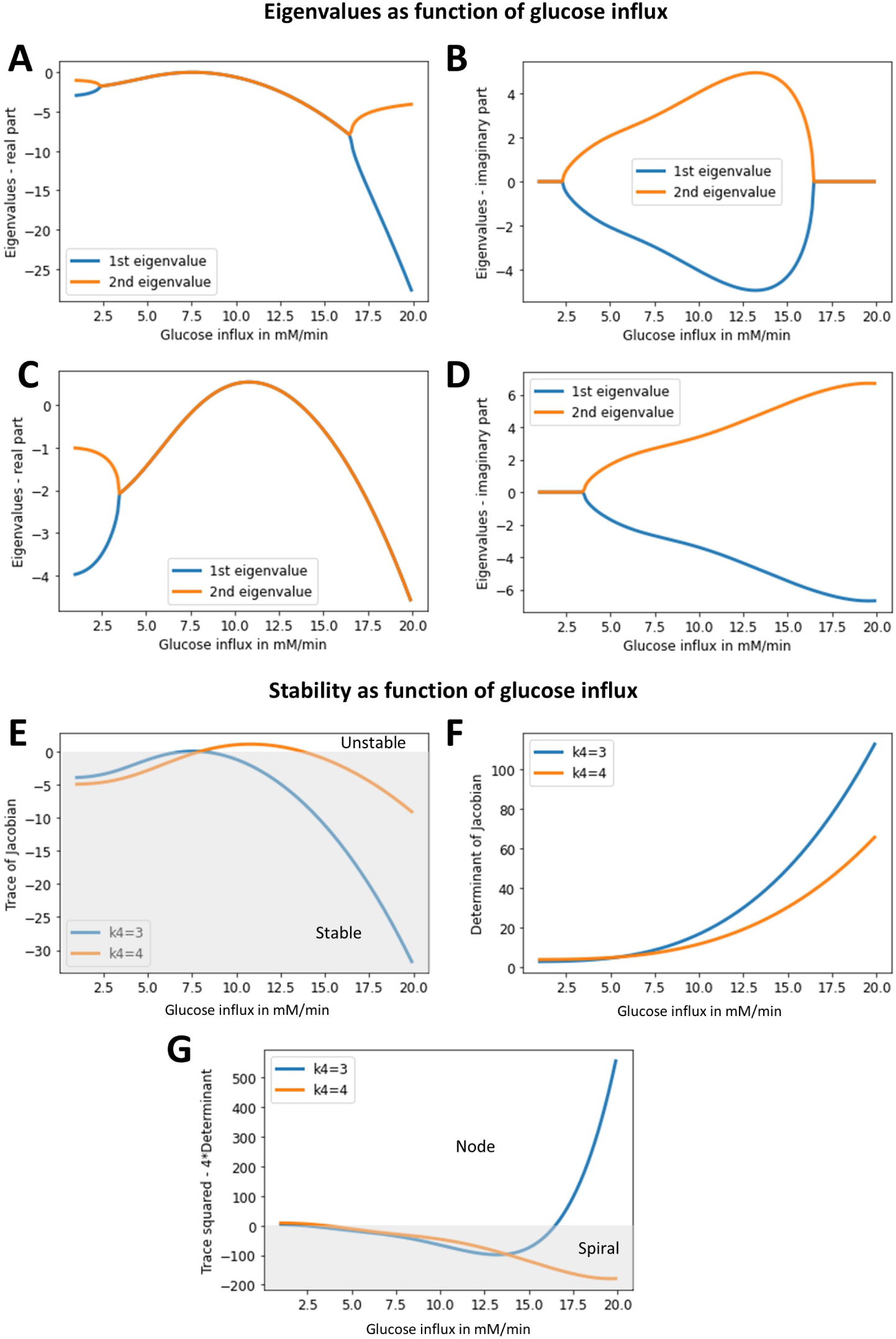
Stability of the steady state determined for the minimal glycolytic model. Based on the calculated steady state values for F6P and F16BP, the eigenvalues of the system were calculated for *k*_2_= 3 min^-1^ (A and B) and *k*_2_= 4 min^-1^ (C and D), and the real part (A and C) as well as imaginary part (B and D) of these eigenvalues are plotted as function of glucose influx, *v*. E-G, a stability analysis was performed as function of glucose influx, showing the trace (E) and determinant of the Jacobian (F) for the rate constant of the aldolase reaction set to *k*_2_=3 min^-1^ (blue lines) or to *k*_2_=4 min^-1^ (orange lines). The sign of the difference between the trace of the Jacobian and four times its determinant informs about the nature of the observed steady state (G). See text and Appendix for further explanations.

### Stochastic variation of glucose influx resulting in damped oscillations can be modeled by DMD

Based on the stability analysis of the derived steady state, we conclude that the oscillatory behavior of the system is very sensitive to the magnitude of the glucose influx, described by the rate *v*, in line with conclusions for the original Higgins model [3]. Sustained oscillations are only found below a critical threshold value, above or below which oscillations become damped and even vanish for high glucose influx (Fig. 1C and D and Fig. 2). Varying glucose influx into glycolysis could be due to stochastic expression of the glucose importers in individual cells or due to varying levels of the hexokinase, which phosphorylates and thereby traps the glucose in cells. This could result in differing levels of damping of the oscillations close to the bifurcation point (Fig. 3A-D). To model this effect, we carried out repeated simulations of the reaction system for stochastically varying influx rate *v* centered around *v* = 14.5 mM/min. In this range of glucose influx varying between ca. 12.5 and 16.5 mM/min a stable focus characterized by damped oscillations will form for most of the simulated cells, while occasionally, i.e., when the glucose influx is around 12.5 mM, also sustained oscillations are found (compare 2C and E with Fig. 3F and G). Drawing 100 samples corresponding to 100 cells from a log-normal distribution of influx rates, as shown in Fig. 3E, confirms this and results in a varying extent of damping of the oscillations for both metabolites (Fig. 3F and G). This clearly supports the central role of glucose influx on the oscillatory behavior of the model, as described previously [3]. Applying DMD to this data matrix after augmentation with time-shifted versions of the simulated time courses results in very accurate reconstruction of the data, even for a low-rank approximation (i.e., r=25) but only as long as the extent of delay embedding is high enough (Fig. 4). For this analysis we constructed the Hankel matrix as described above and in the Materials and methods section, i.e., we stacked *d* shifted versions of each row (resembling a single time series for one value of the glucose influx) consecutively and applied standard DMD to this augmented matrix. A systematic analysis of the extent of time-delay embedding reveals that the eigenvalue spectrum becomes stable for delays *d* > 120, coinciding with good reconstruction quality for the damped oscillations for each cell (Fig. 4C-G). In fact, the reconstruction quality converges towards low RMSE values above *d* = 150 (Fig. 4G). For comparison, we calculated the analytical eigenvalues for the same data set, rescaled them to the unit circle and plotted them on top of the eigenvalue spectrum for d=150 (Fig. 4c, last panel, yellow and black dots).

**Figure 3.**
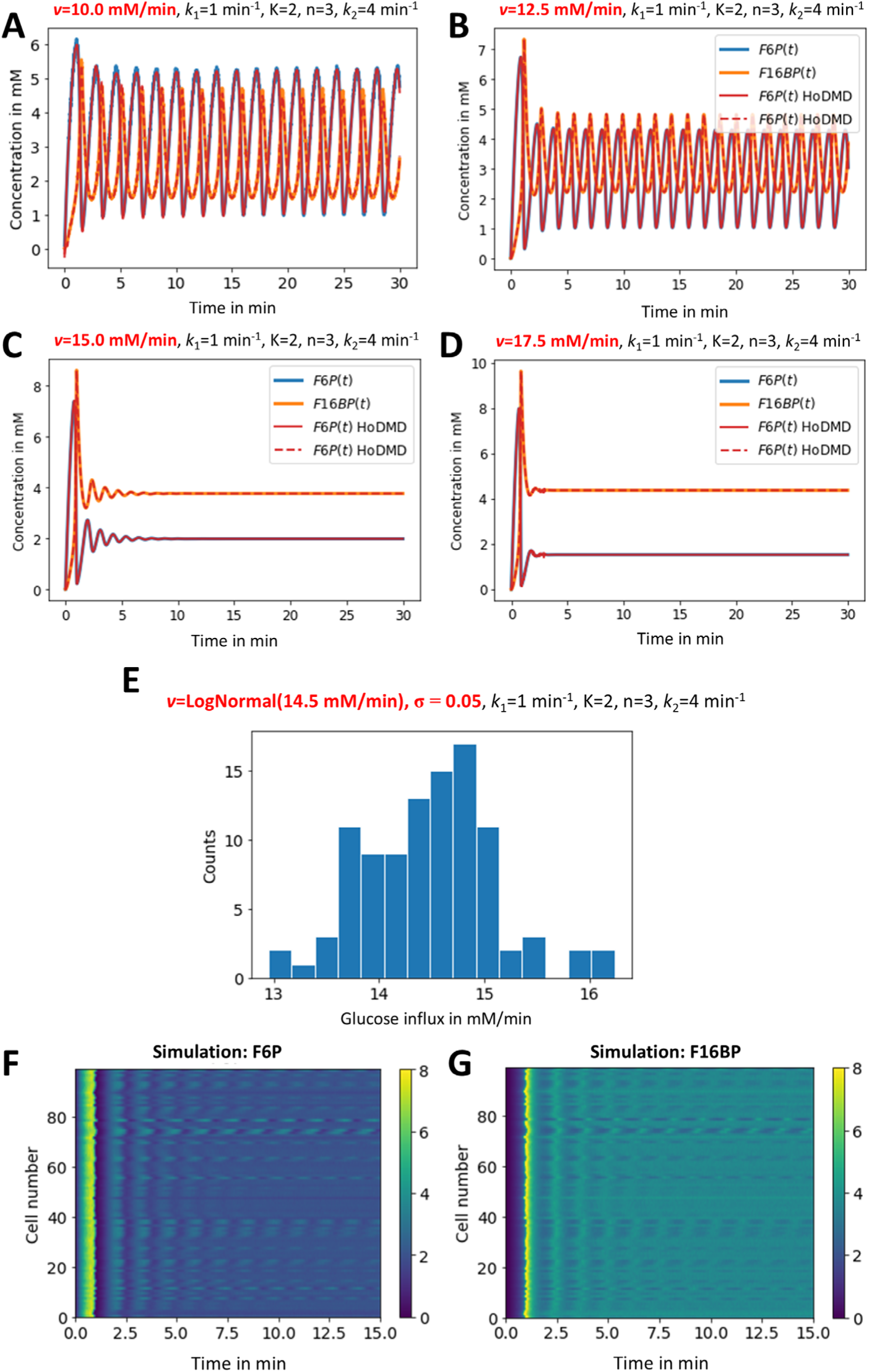
Higher-order DMD models glycolytic oscillations as function of glucose influx. Time courses for F6P and F16BP were simulated with the minimal glycolysis model for varying glucose influx described by the rate v (red values on top of panels A-D, blue and orange dots show simulation of F6P and F16BP, respectively, while red straight and dashed lines show the corresponding HoDMD reconstructions). A distribution of 100 influx values was generated from a log-normal distribution centered around 14.5 mM/min (E). Simulation of time courses for F6P (F) and F16BP (G) using these influx values and the other indicated parameters (see top of panel F and G).

**Figure 4.**
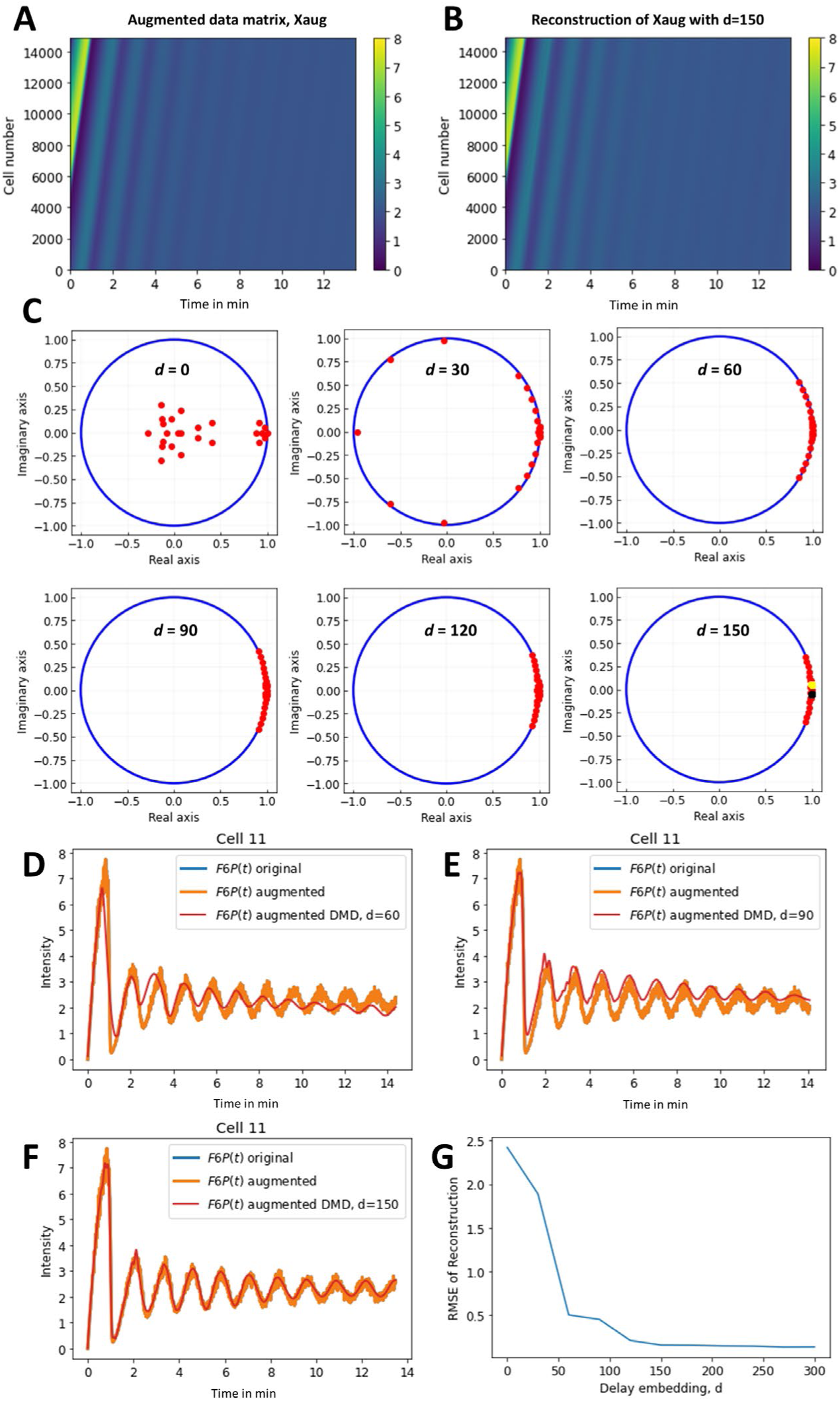
Generation of Hankel matrix and its analysis using DMD for varying delays. Time-delay embedding was realized by generating Hankel matrices of the simulations of the first species, i.e., F6P, as shown in Fig. 3F. For that, time-shifted versions of each simulated oscillation were generated and stacked as additional rows to the original data matrix, giving an augmented data matrix (A). Applying DMD to this augmented matrix results in very good reconstructions (B). Eigenvalue spectrum of the identified DMD modes for varying extent of delay embedding (C). For increasing *d*, the eigenvalues shift to the right and concentrate around the ones determined for the simulated system (yellow and black dots on plot for *d* = 150, see Eq. A8 and A9). D-F, simulated time courses for a selected cell (Cell 11), with original data from matrix X in blue, data from the augmented matrix Xaug in orange (which is identical to those of matrix X)

This comparison shows that the eigenvalues determined by DMD concentrate around the analytical eigenvalues for increasing delay embedding, while at the same time the reconstruction quality increases. Since the analytical eigenvalues are derived from the system linearized around the steady state (see Appendix), this important result confirms that DMD achieves an optimal linear approximation of the non-linear dynamic system [17, 45]. Note that there are 200 analytical eigenvalues for the entire simulated data set of 100 cells, i.e. 2 conjugated eigenvalues for each cell. In contrast, the DMD approximation only contains 25 eigenvalues for the entire data set due to the low-rank truncation of the Koopman operator with the time-delayed snapshots as observables [37]. In the reconstruction, each time course is represented by a weighted sum of these 25 eigenvalues and eigenfunctions (see Eq. 10 and 11 in Materials and methods). Thus, while the state dimension of the input data is increased by delay embedding, the identified dynamic modes span a low-dimensional subspace of the infinite-dimensional Koopman operator, allowing for faithful reconstruction of the observed dynamics. This is nicely illustrated, when looking at the attractor of the simulated system in comparison to the attractor reconstructed by DMD with time-delay embedding (Fig. 5). Both the attractors of a stable focus and of limit cycles caused by differences in glucose influx for individual cells are correctly reconstructed by DMD (Fig. 5A-D). Moreover, the reconstructed manifold *M*_DMD_ is much smoother than the simulated one, emphasizing the potential of DMD for time series denoising. It has been suggested that changing the delay step length can improve reconstruction quality of the embedding [46], so we tested this by increasing the step length for the delay embedding from 1 to 5, but this gave almost identical results (Fig. 5E and F). Together, DMD with time-delay embedding is a powerful tool to dissect and reconstruct complex metabolic time series of non-linear biochemical models using a linear model in higher dimensions.

**Figure 5.**
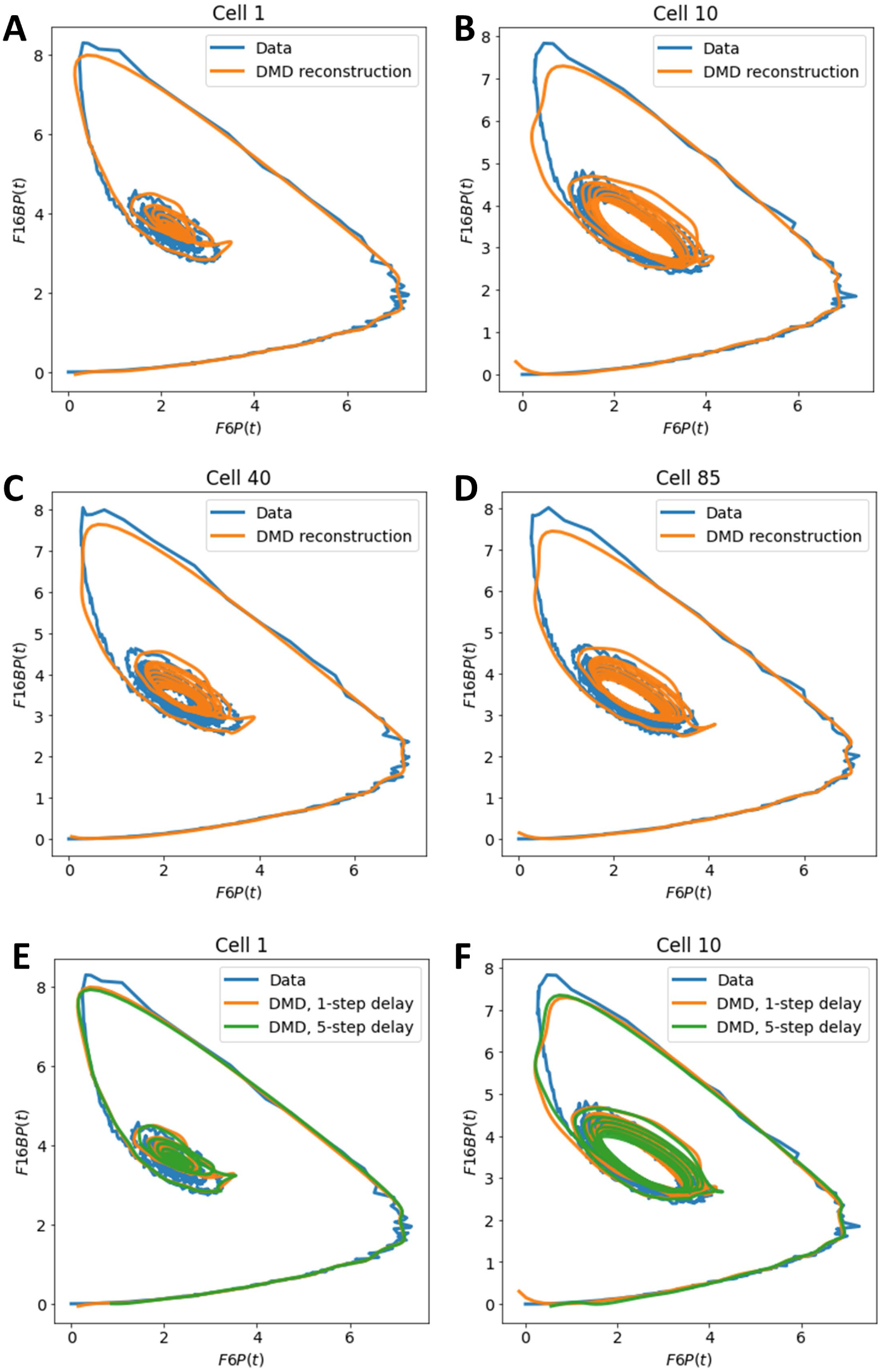
Reconstruction of the system attractor using DMD with time-delay embedding. The two-state glycolysis model was simulated with added noise and analyzed using DMD as described in the legends to Figs. 3 and 4. The time course of the two metabolites, F6P and F16BP, was plotted against each other for selected cells of the simulation with stochastic glucose input, which are indicated on top of each panel (blue lines). The corresponding reconstructions of the time courses are plotted for a 1-step delay in orange and for a 5-step delay in blue for the same cells. Cell 1 is shown in (A) and (E), cell 10 in (B) and (F), cell 40 in (C) and cell 85 in (D).

### Analysis of starvation-induced oscillations of NADH autofluorescence in yeast cells by DMD

To assess the performance of DMD with delay embedding in reconstructing real metabolic oscillations, we recorded cellular autofluorescence in baker’s yeast Saccharomyces cerevisiae. The intrinsic fluorescence of NADH is a measure of glycolytic flux and has been studied extensively in yeast and other eukaryotic cells (see Introduction). We used a yeast strain, which is unable to rely on oxidative phosphorylation due to defective heme synthesis (i.e., *hem1*Δ cells). Without pre-starvation, these cells only show a transient increase in NADH fluorescence upon glucose addition but no or only very little subsequent oscillations (Fig. S3A-C). In contrast, when being pre-starved, the cells respond to addition of glucose with (damped) oscillations of NADH fluorescence (Fig. 6). Time traces of autofluorescence were extracted as mean intensity per cell after segmenting individual cells. For segmentation, we have trained a deep-learning model, and the mean intensity per cell was afterwards aligned in a cell-time plot, exactly as for the simulations (see Materials and methods and Fig. 6A). DMD with time-delay embedding but not standard DMD reconstructs the time series for each cell accurately, as long as the extent of embedding is sufficient (i.e, *d* = 150; Fig. 6B and C). These results agree with our simulation results (see Fig. 4), and they are also supported by earlier findings showing that the extent of delay embedding needed for faithful reconstruction of time series depends on the spatial dimension of the data [26, 27, 29]. In fact, as shown by Mezic and co-workers, the true Koopman eigenvalues and eigenfunctions for an ergodic system can be found from a Hankel matrix in the limit of infinite-time observations [37]. Since the experimental metabolic oscillations are not stationary, it is interesting, that we still achieve very good approximation of the data using DMD with time delays.

**Figure 6.**
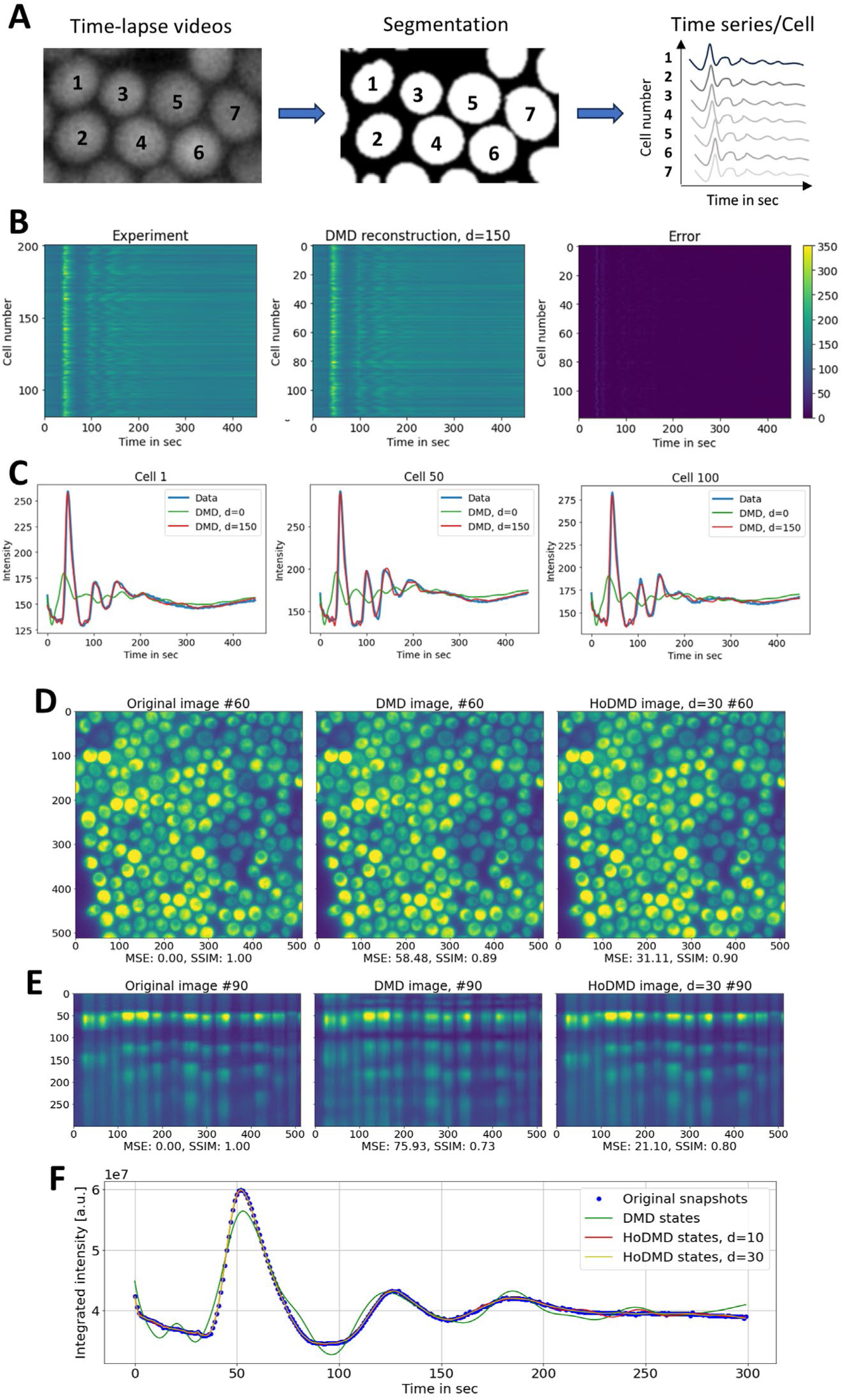
DMD reconstruction of experimental autofluorescence oscillations in yeast cells. Yeast cells were pre-starved, washed and placed on a microscope dish coated with poly-D-lysine before imaging on a wide field microscope as described in Materials and methods. A-C, video recordings were segmented and individual time traces of mean intensities arranged as matrix, exactly as used in the simulations. B, experimental data sets (left panel) were reconstructed using DMD with time-delay embedding of *d* = 150 (middle panel), resulting in a very low reconstruction error (right panel). C, intensity traces of selected cells from the experimental data (cell number is given on top of each panel) are plotted (blue lines) and compared to the DMD reconstruction without embedding (*d* = 0, green lines) and with embedding (*d* = 150, red lines). D-F, HoDMD reconstruction of entire video sequence without segmentation of individual cells. D, original time-lapse recording (left panel), DMD reconstruction (without delay embedding, middle panel) and HoDMD reconstruction (with *d* = 30, right panel), all shown for frame #60 of the corresponding image stack. E, the same as D but side view (xt-view) along the horizontal line #90. Image quality measures, i.e., MSE and SSIM, relative to the original data are given below each panel. F, integrated intensity along the time axis for raw data (blue dots), DMD reconstruction with *d* = 0 (green line), HoDMD with *d* = 10 (red line) and *d* = 30 (yellow line), respectively.

A potential problem of the above analysis is, that a finite-dimensional approximation of the infinite-dimensional Koopman operator can result in spectral pollution, i.e., the occurrence of spurious eigenvalues that are not related to the evolution operator [47]. While the Hankel-matrix based DMD method theoretically ensures to recover the correct spectrum of the Koopman operator (see above), this is only guaranteed in the limit of infinite observations for an ergodic system [37]. Also, the Koopman operator can have a continuous spectrum, particularly for chaotic systems [47]. To validate the eigenvalues identified by DMD with delay embedding we re-analyzed the simulated and experimental data sets using residual DMD, a method to identify pseudo eigenfunctions based on calculations of residuals for each eigenvalue pair [47, 48]. We find for both the simulated and the experimental data, that without delay embedding, a portion of the eigenvalues, particularly towards the center of the unit circle, are associated with large residuals, indicating that they are spurious eigenvalues (Fig. S4, left columns) [48].

In contrast, with delay embedding (*d* = 150), a larger part of the eigenvalue spectrum is associated with low residuals showing that more of the calculated basis functions actually contribute to a linearized description of the data. As a result, we achieve a better approximation of the Koopman operator and thereby of the simulated and measured fluorescence dynamics, respectively (Fig. S4, right columns). This result further emphasizes the power of delay embedding as part of the DMD method in dissecting non-linear dynamical systems [26]

To analyze, not individual cells, but the entire video data at once without prior cell segmentation, we returned to the HoDMD analysis (compare Fig. 1, 3, 6D-F and S6). This is because HoDMD has been shown to be particularly useful for analyzing multi-dimensional time series by taking the history of the data into account [25]. Again, HoDMD but not standard DMD can reconstruct the transient oscillations found in starved yeast cells upon addition of glucose, while standard DMD without time-delay embedding was not successful (Fig 6D-F). A higher time-delay embedding was required for reconstructing the dynamics of non-starved cells compared to starved cells (compare Fig. 6D-F with Fig. S3). This is likely a consequence of the more transient nature of the dynamics in non-starved cells and not related to cross differences in the singular value spectrum of both data sets (Fig. S7). Indeed, it has been shown previously that time series with transients decaying faster than the slowest mode represent challenges for DMD [49]. For both situations, i.e., the autofluorescence dynamics of starved and non-starved cells, the singular values of the snapshot matrix generated from the time-lapse video sequence decay rapidly (Fig. S5). This low-rank structure of the data is a pre-condition for the DMD reconstruction with few modes governing the essential dynamics of the system.

### Prediction of metabolic oscillations by DMD with time-delay embedding

To assess the ability of DMD to predict metabolic oscillations, we truncated the simulated or experimental time series at selected time points and assessed the ability of DMD to reconstruct the training data and to predict unseen data. We found that DMD with time-delay embedding allows for predicting damped oscillations for several periods in the future but also, that predictions become less accurate, the more sustained the oscillations are (Fig. S6A-D). To assess the prediction accuracy of DMD in the critical parameter range, in which the system forms a limit cycle, the simulations were repeated for a mean glucose influx of v = 9.5 mM/min with σ = 0.05 mM/min. For the resulting sustained oscillations, DMD’s ability to predict future time points is worse but can be improved by increasing the delay embedding dimension, *d* (Fig. S6E-H). The limited capability of DMD to predict future time points in highly non-linear systems is also the outcome of other studies and a consequence of the underlying assumptions, the approximation of non-linear dynamics by a linear and finite matrix *A* [50].

Based on these findings, we assessed the ability of DMD with time-delay embedding to predict future time points of the experimental autofluorescence time series. For that, experimental data were averaged and split to create the dataset used for training and prediction. Various truncation and delay embedding steps were systematically tested to identify the optimal configuration for capturing the dynamics of the system. Not unexpectedly, DMD performs better with respect to predicting future time points, the more data it has seen, i.e., the deviation from real unseen data becomes smaller for more data points in the training set (Fig. 7). When only the first 100 data points were in the training set, DMD predictions were very poor with very negative R^2^-values (< 400) indicating that the residual error is much larger than the data variance itself. Also for 200-400 data points out of the 600 total in the training set, the R^2^-values were negative, but they approached R^2^=0, which corresponds to the model being as good as the data mean for prediction. With respect to the training data, using a delay embedding of *d* = 25 corresponding to 25% of the truncated data for the 100 data points does not fit the training data well and fails to predict future time points (Fig. S7A). However, when the delay step is increased to 50% of the data points, the model achieves a near-perfect reconstruction of the training data, even for this low number of seen data points, while it still fails on the test data (Fig. S7B). Conversely, increasing the delay step further to 75% of the 100 data points in the training set results in a poor fit once again in both, the training and test data (Fig. S7C). This trend is also seen for truncating the time series after 400 time points (Fig. S7D-F) and is, in fact, consistent across all truncation scenarios (see Fig. S7G-H). The quality of the reconstruction and prediction of the time series by DMD are quantified in the R^2^ plot shown in Fig. S7G for all data (i.e. training and unseen test data) and in Fig. S7H for the training data, only. The reconstruction performance is clearly highest for 50% delay embedding, and the difference to 25 and 75% delay embedding grows, the fewer data points enter the training (Fig. S7H). The ability to predict future time points was also assessed for HoDMD on entire video sequences. For that, the HoDMD model was trained with 150 out of the 300 image frames.

**Figure 7.**
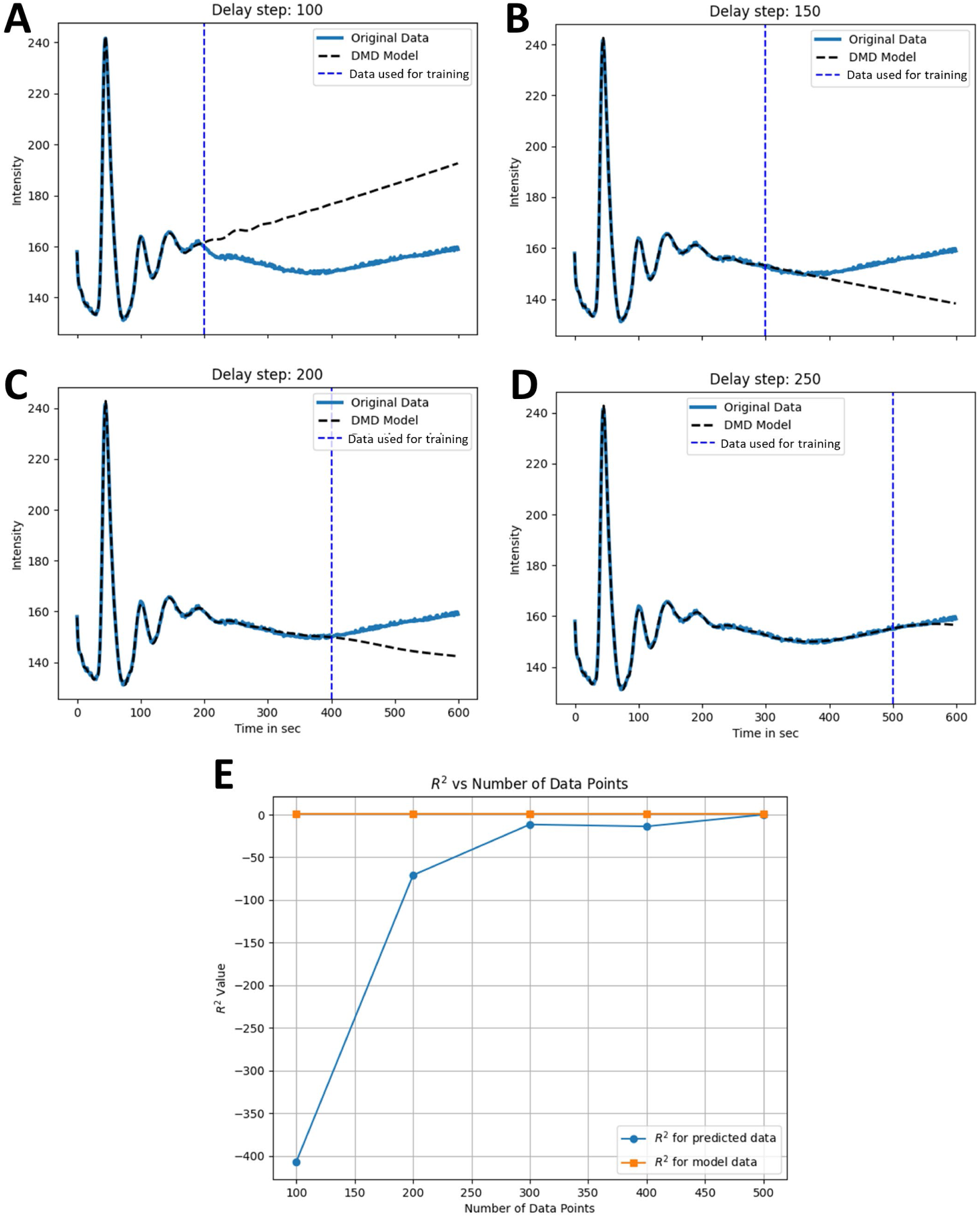
Prediction of experimental time courses using DMD with time-delay embedding. Experimental time courses of NADH fluctuations recorded in response to addition of 15 mM glucose were averaged for all cells in a field (blue lines in A-D, labeled ‘Original data’) and reconstructed using DMD with time-delay embedding (black dashed lines in A-D, labeled ‘DMD model’). Training data started at t=0, and its length was varied with 200 time points in A, 300 in B, 400 in C and 500 time points in D, respectively. This is indicated by the dashed blue line in panel A-D (labeled ‘Data used for training’). E, R^2^-values as function of data points in the training set for the training data (‘model data’, orange line and symbols) and the test data (‘predicted data’, blue line and symbols). See text for further explanations.

Despite accurately reconstructing the phase-shifted oscillations of individual yeast cells, HoDMD cannot predict the damped oscillations of cellular autofluorescence apart from the first few frames (Fig. S8). We conclude that the ability of DMD to predict new data points for non-linear time series of glycolytic oscillations is very limited, even after time-delay embedding. To compare the prediction capability of DMD with delay embedding with other architectures, we re-analyzed the experimental autofluorescence oscillations with an LSTM, a form of recurrent neural networks (RNNs). Compared to classical RNNs, LSTMs are designed to prevent exploding or vanishing gradients, when learning long-term dependencies in the time series data [51]. This is achieved by implementing a unique architecture consisting of an input gate into the memory cell, the basic unit of an LSTM, an output gate and a forget gate, which removes unnecessary information from the network at each training step. LSTMs are designed for forecasting, as they capture temporal dependencies over long time horizons. To evaluate the performance of the model, the training dataset was prepared by sequencing the data into overlapping segments, akin to delay embedding. For example, one segment might include elements 0 to 20, while the subsequent segment spans elements 1 to 21, and so on. This approach ensures the preservation of temporal dependencies within the data. Once the model was trained, predictions for the remainder of the dataset were conducted. Starting with the last value of the training set, the model was used to predict the subsequent value. This predicted value was then iteratively fed back into the model to generate the next value, allowing for sequentialprediction of the entire dataset.When applying an LSTM with only one layer to the experimental data with a training time of the first 100 snapshots, the ability to predict future time points was poor and only slightly increased by extending the sequence length (Fig. S10A and B). Including more layers increased the prediction accuracy, but only up to 6 layers, beyound which the R^2^ became worse again, likely due to overfitting. We conclude that the overall performance of DMD with delay embedding and LSTMs to reconstruct and predict the experimental data is comparably low.

## Discussion

Metabolic oscillations are a widespread phenomenon and occur not only in glycolysis of fermenting yeast cells but also in glycolysis of cancer cells, in pancreatic β-cells in the connection with pulsatile insulin secretion, as well as in muscle cells and human neutrophils [1, 34]. A lot of theoretical work has been dedicated towards obtaining a mechanistic understanding of such oscillations, but little work has been directed towards employing modern data-driven methods for quantitative analysis of such oscillations based on real time-lapse microscopy data of individual cells [13, 16]. Here, we show that DMD combined with time-delay embedding allows for faithful reconstruction and modal decomposition of simulated and experimental glycolytic oscillations. We show that the eigenvalue spectrum determined by DMD for simulated and experimental metabolic oscillations depends on the delay embedding dimension. For increasing delays, the spectrum converges and becomes stable with low residuals, showing that under those conditions, DMD recovers a good linear approximation of the Koopman operator, capturing essential elements of the system dynamics. The embedding dimension plays a central role in approximating the dynamic manifold for the oscillations, which is a result of Taken’s theorem [26]. We demonstrate, that this is even the case for non-stationary dynamics and in the presence of noise, as long as the system is relatively close to a stable steady state. However, we also find that DMD’s ability to predict future time points is limited for both, the simulated and experimental glycolytic oscillations. The limited performance in predicting future time points is a direct consequence of the assumptions underlying any DMD analysis, namely that the dynamics are approximately linear. More precisely, DMD assumes that there exists a linear subspace of the Koopman operator, which captures the essential system dynamics as a linear superposition of exponential eigenfunctions with constant amplitudes and eigenvalues. The validity of this assumption, however, is questionable in situations, where the system quickly moves away from the fixed point, i.e., in which the Jacobian defining the tangent space poorly describes the system dynamics (see Appendix) [45]. In these situations, higher-order terms of the Taylor expansion around the steady state play an important role, and the linearity assumption becomes questionable, particularly in the presence of noise. This effect will be even more pronounced for the experimental situation. While inclusion of higher-order terms, as in HoDMD as well as time-delay embedding increase the state space, these DMD variants are still linear methods, as they attempt to find a finite-dimensional approximation to the (linear) Koopman operator. Thus, the DMD approach does not explicitly model the non-linear terms that characterize the system’s true behavior. Instead, the inclusion of higher derivatives, as in HoDMD, or the use of time-shifted versions of the data as in DMD with time-delay embedding, leverage additional information about the system to refine the linear approximation.

LSTMs, on the other hand, are designed to model complex statistical dependencies in the time series data, making them well-suited to model and predict non-linear and non-stationary metabolic oscillations. This is partially a consequence of the use of non-linear activation functions in their architecture but also due to the underlying memory mechanism in such networks, which allows for dynamic adaptation to non-linear variations in the time series data [51]. These properties provide LSTMs with the power to dissect complex time series into multiple patterns happening on different scales, e.g. fast oscillations superimposed on slower trends. The idea of including past information into reconstructing the current state and in forecasting future time points is common to both LSTMs and time-delay embedding. LSTMs have the ability to adapt their memory length via the gating mechanisms, giving the latter much more flexibility in handling non-stationary and non-linear dynamics. On the other hand, LSTMs learn only an implicit representation of the attractor describing metabolic oscillations, while time-delay methods explicitly reconstruct this attractor. Based on these differences we would have expected that LSTM is much more powerful than DMD to predict future time points of the metabolic oscillations. However, we found that the performance of both methods is comparable and in general rather poor, likely because both methods are equally challenged by the non-stationarity of the experimental data, in which no repeating pattern can be predicted.

Using time-delay embedding as the coordinate basis for data-driven representation of the Koopman operator is powerful, as shown here and in many other applications including analysis of oscillating signals in neuroscience experiments [41] [52], and of 3D or polarization fluorescence microscopy data [43]. To further improve the performance of this approach, one could explore non-uniform embedding strategies, as reviewed and suggested in [46]. One could also explore alternative observables, for example, use a dictionary of non-linear functions of the snapshot data as observables, as suggested in various forms of extended DMD [37, 53]. By using such observables, one would eventually increase the likelihood to find good eigenfunctions of the Koopman operator, which adequately describe the entire manifold. However, finding such observables is not trivial, and a lot of current efforts are dedicated towards this goal [54, 55]. One possibility is to combine DMD with neural-network based approaches in the future. This would have the aim to learn a faithful linear encoding of the non-linear dynamics allowing for improved finite-dimensional approximation of the Koopman operator [53, 55–62]. Alternatively, the DMD approach could be informed about forcing terms, which exert control of the studied dynamic system. This approach is also widely used, and when combined with delay embedding, it was found to describe even chaotic systems [29, 63]. Finally, the assumption of eigenfunctions being constant over the entire time series can be replaced by dynamic modes varying along the studied dynamics. This has been implemented as multiscale reconstruction, similar to wavelet analysis [18], and as windowed approach in the recently developed non-stationary DMD [64]. Like some of the autoencoder-based DMD variants [60], these approaches might be better suited to analyze non-stationary oscillations, and they will be explored for imaging-based analysis of the dynamic metabolism of living cells in the future.

## Materials and methods

### The standard DMD algorithm applied to simulated and experimental time courses

The theory behind DMD and HoDMD is described in [25, 65], and follows the procedure outlined in our recent publications [23, 24, 43]. Briefly, snapshots in time (i.e., concentration of metabolite species or fluorescence intensities of NADH, either as single measurements or as images) are arranged in a sequence [22, 66]:

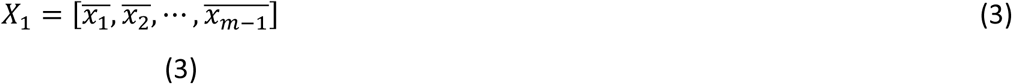

and

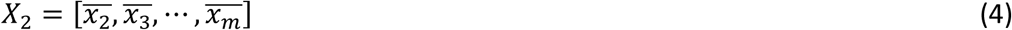

The index *k*=1,…, *m* runs over all acquired snapshots in time with steps, Δt. One can define the progression from state x(k·Δt)=x_k_ to x_k+1_ as:

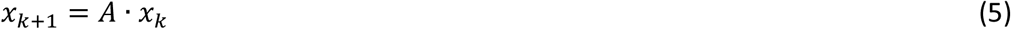

Here, *A* is a matrix which describes the advancement of the system from image *x*_k_ to image *x*_k+1_. This matrix resembles the Koopman or transfer operator for measurements g(*x*_k_)=*x*_k_ [65]. With that the system corresponding to Eq. (5) becomes *X*_2_ = *A*·*X_1_*, from which we can find *A* by minimizing the Frobenius norm, ‖·‖_*F*_ [65]:

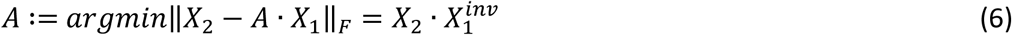

To solve this system, one first finds the pseudoinverse of the first data matrix, 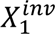. For that, one uses a SVD of *X*_1_ into unitary matrices U ∈ *R*^n·(m-1)^, V*∈ *R*^n·n^ with singular values in the diagonal matrix Σ ∈ *R*^n·(m-1)^:

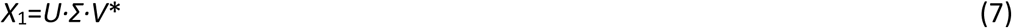

To obtain a low-rank approximation of the dynamic only the modes up to rank *r* < (*m*-1) are retained, i.e., one approximates the system matrix *A* by projecting it onto the left singular vectors, which are the column vectors of U. This gives a much smaller matrix A’ of maximal size *m* times *m* via a similarity transformation [20, 65]:

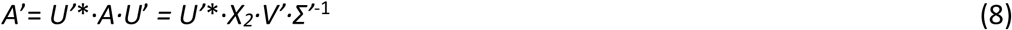

Here, *U*’, *V*’ and *Σ*’ are rank *r* approximations of the full matrices, *U*, *V* and *Σ*, and the complex conjugate transposed matrix is indicated by *:

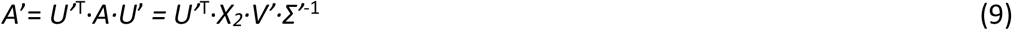

The similarity transformation of Eq. 8 corresponds to a dimension reduction, which reduces the size of the system matrix from *A* ∈ *R*^(m-1)·(m-1)^ to *A’*∈ *R*^r·r^ giving a low-dimensional representation of the dynamic system [65]. To obtain the spectral decomposition of the reduced system, one finds the eigenvalues, *λ_j_* and eigenfunctions, *φ_j_*, for each DMD mode *j*, by solving the corresponding eigenvalue problem:

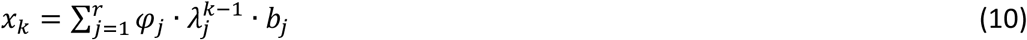

For a continuous system, e.g. in time, one can rescale the eigenvalues according to *ω* = ln(*λ*/Δ*t*), such that Eq. (9) can be written as [19]:

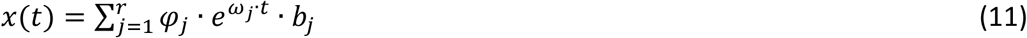

Here, *x*(*t*) is a vector of images (x, y index omitted for brevity) as a function of time, *t*,

#### Extension of the DMD algorithm to include time-shifted versions of the snapshot data

The matrix *A* can be approximated from the data by first introducing the higher-order Koopman assumption, which reads

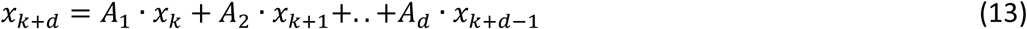

Here, *k* runs from 1 to *m*-*d*, where *m* is the number of snapshots taken at discrete time steps and *d* is the set time-delay. A rank-truncated SVD, as described in Eqs. 7 to 9 is applied to the snapshot matrix leading to:

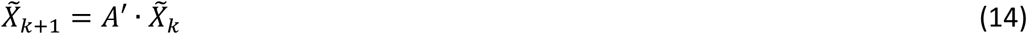

The reduced matrices for the snapshots and the evolution of the system read [25]:

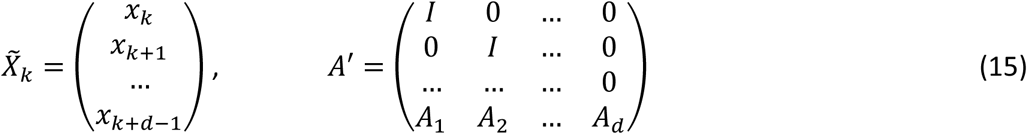

HoDMD uses this approach and was employed on the simulated and image data as implemented in the PyDMD package [42]. An alternative approach for time-delay embedding is to construct a Hankel matrix of the data matrix of Eq. 3 by including time-shifted versions of each row according to:

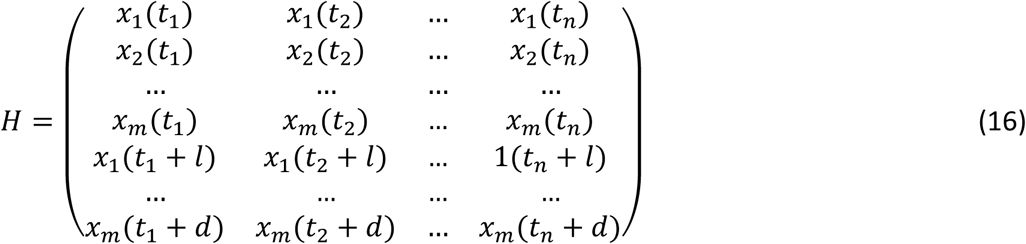

Here, *t*_i_, *i* = 1, …*n* are the snapshots and *x*_j_ = 1, …*m* are the individual time traces, i.e., the simulated or experimental time courses for each cell. The index *l* is the time delay (chosen as *l* = 1 or 5) and *d* is the embedding dimension, i.e., the total number of chosen time shifts. The matrix H and its time-advanced variant were used in the standard DMD algorithm according to Eqs. 3 to 11, above.

### Yeast cell culture, live-cell imaging and cell segmentation

BY4741 cells were grown in YPD media consisting of 2% (w/v) D(+)-Glucose Monohydrate (Merck, 1.08342.1000), 2% (w/v) Bacto Peptone (BD Chemicals, 211,677), 1% (w/v) Yeast Extract (Merck, 1.03753.0500) and 0.02% (w/v) adenine (Sigma-Aldrich, A-2786). The yeast strain *hem1*Δ is derived from W303-1*α* (MAT*α*ade2-1 his3-11,15 leu2-3,112 trp1-1 ura3-1 can1-100), and it was kindly made available by Dr. Thomas Pomorski (University of Copenhagen, Section Transport Biology). *hem1*Δ cells were grown in YPD media with 50 µg/ml δ-aminolevulinic acid (dALA) (Sigma-Aldrich, A3785).

Before imaging, the media was changed to SD media without glucose consisting of 0.001% (w/v) D(+)-Glucose Monohydrate (Merck, 1.08342.1000), 0.7% (w/v) Yeast Nitrogen Base w/o amino acids (BD Difco, 291920), 0.2% (w/v) Yeast Synthetic Drop-Out Medium Supplements without uracil (Sigma-Aldrich, Y1501), 0.012 % (w/v) adenine (Sigma-Aldrich, A-2786) and 0.002% (w/v) uracil (Sigma-Aldrich, U1128) and incubated in a shaking incubator (∼30°C, 150 rpm) for 3 h to ensure glucose depletion. The yeast was diluted to a concentration of 1 or 2 OD_600_/ml in PBS before the cells were added to a microscope dish. During imaging, glucose was added to the dish to a final concentration of 15 or 30 mM.

Imaging was conducted on a Leica DMIRBE fluorescence microscope with a 100 x 1.3 NA oil immersion objective. The NAD(P)H autofluorescence was obtained using an A4 filter cube with a 360 nm (40 nm bandpass) excitation filter, 400 nm dichromatic mirror and a 470 nm (40 nm bandpass) emission filter.

Each cell was segmented using a Cellpose model on the max Z-projection of the fluorescence image. Based on individual cells, the mean fluorescence over time was extracted. Acquired images were segmented using the deep learning-based segmentation method Cellpose, as described [67]. The brightfield images were segmented using the built-in CPX model with a diameter of 40 in CellPose.

The LSTM analysis was conducted on the averaged experimental data. The LSTM model was implemented using PyTorch, with mean squared error as the loss function and the Adam algorithm as the optimizer. Once the model was set up, the parameters to be tested were defined, including the number of layers and nodes in the network, as well as the proportion of data used for training. After defining these parameters, the training process was initiated. The network was trained until the loss function no longer decreased, typically after 100–200 epochs. For analysis, the model’s predicted data was plotted against the real, unseen experimental data, and a quantitative evaluation was performed by calculating the r² value.

## Supporting information

Supplemental data refered to in the manuscript.

## Data availability

The datasets used and analysed during the current study are available from the corresponding author on reasonable request.

## Acknowledgements

DW acknowledges funding by the Danish Research Council (grant ID: 2032-00139B). Special thanks to Dr. Matthew Colbrook from Oxford University for providing the code for the Residual DMD and for helpful discussions. We are also acknowledging Line Lauritsen for technical assistence and for critical reading of the manuscript.

## Appendix

### Stability analysis of steady state of the glycolysis oscillation model

The steady state concentrations of F6P, 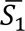, and of F16BP, 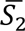, read:

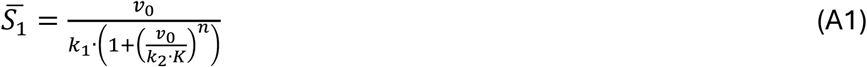

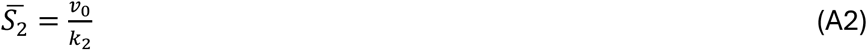

The stability of the steady state can be determined from the Jacobian matrix, which approximates the non-linear dynamics by the first term of a Taylor-expansion according to:

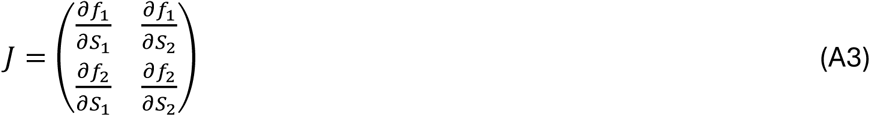

Here, *f*_1_ and *f*_2_ correspond to the two differential equations given in Eq. 1A, B, and the four matrix entries read:

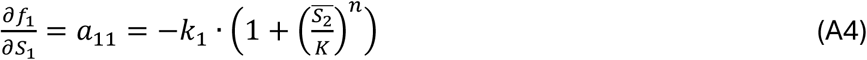

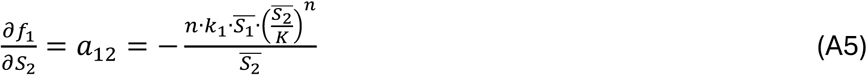

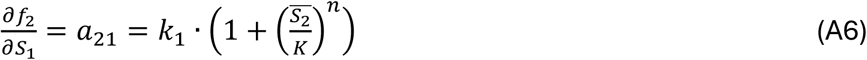

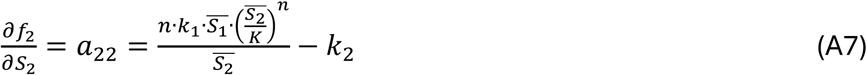

The eigenvalues of the Jacobian matrix read:

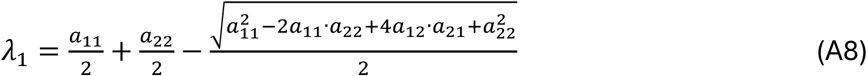

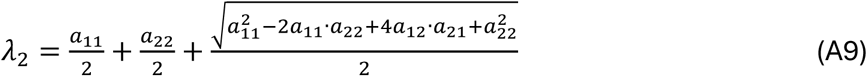

Since all parameters (i.e., inflow rate, *v*_0_, rate constants, *k*_1_ and *k*_2_, equilibrium constant, *K*, and Hill coefficient, *n*) are larger than zero, one can immediately see that always *a*_11_ < 0 and *a*_12_ < 0 and *a*_21_ > 0. The last element, *a*_22_, can be positive or negative, depending on the value of the rate constant *k*_2_ compared to the first term in Eq. A7. The eigenvalues can be real or complex, as further explained in the main text. The trace of the Jacobian is

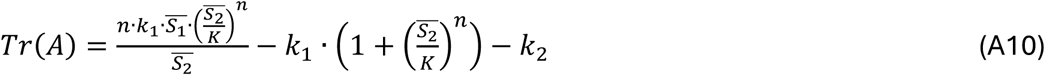

And its determinant reads

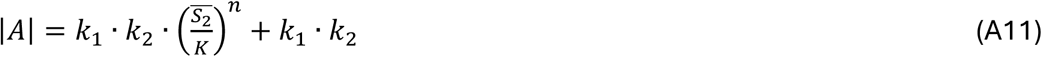

One sees that |*A*|>0, always. Together with the eigenvalues, the trace and determinant, the roots of the characteristic equation of the Jacobian can be calculated which, following the Hurwitz criterion, inform about the stability of the steady state (see main text).

