## Supplemental data refered to in the manuscript. for "Dynamic mode decomposition for analysis and prediction of metabolic oscillations from time-lapse imaging of cellular autofluorescence"

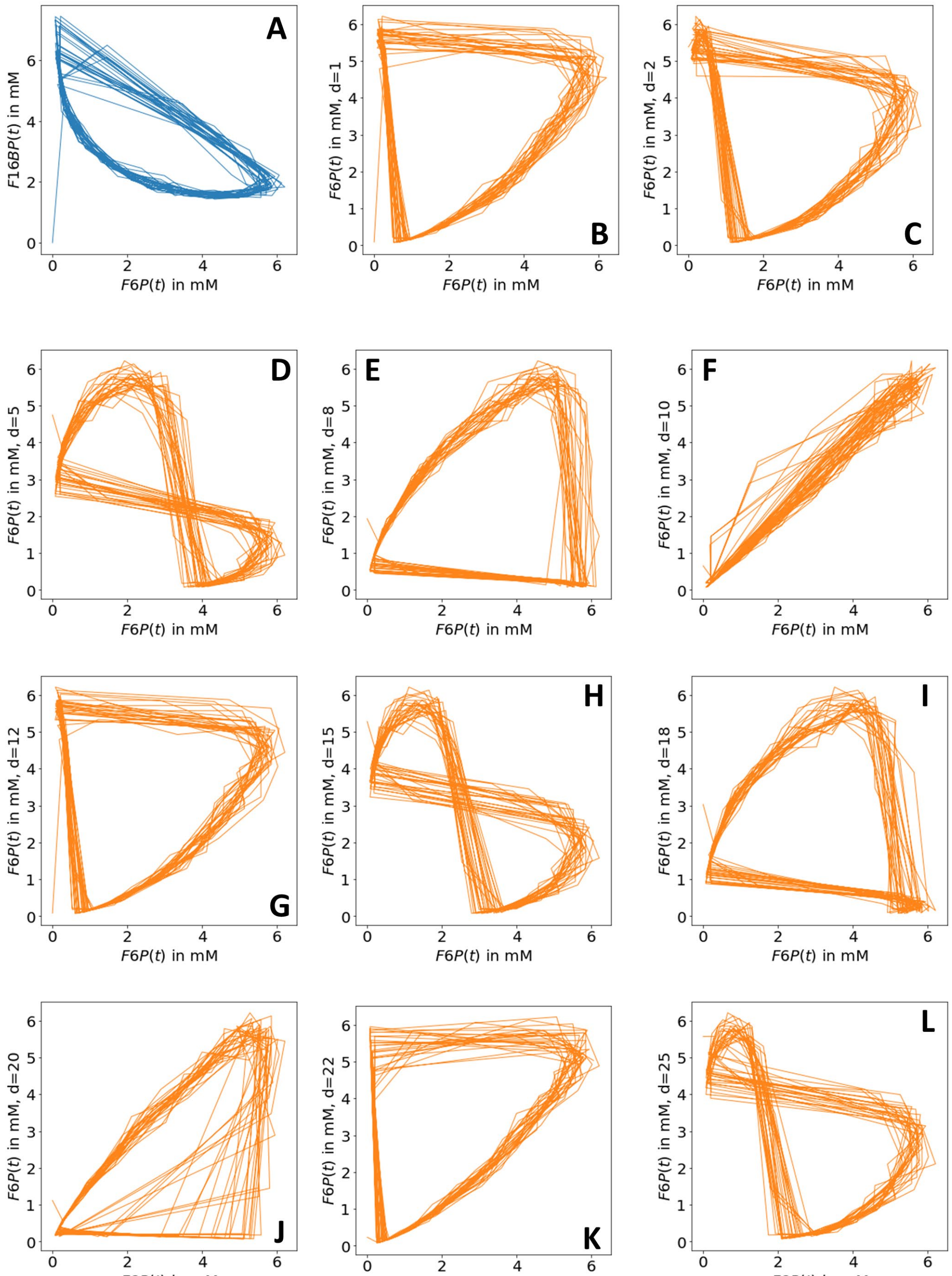


**Figure S1. Time-delay embedding can approximate the attractor of the glycolysis model.** The glycolysis model was simulated for the parameters v=9.5 mM/min, $k_1$=1 $s^{-1}$, K=2, n=4 and $k_2$=4 $s^{-1}$ with added log-normal noise, as described in Fig. 1 and 3 of the main text. A, phase plot of time course for F16BP plotted against that of F6P. B-L, time course of F6P plotted as function of time-shifted versions of itself for increasing time-shift (delay, *d*); i.e., for *d*=1 (B), 2 (C), 5 (D), 8 (E), 10 (F), 12 (G), 15 (H), 18 (I), 20 (J), 22 (K) and 25 (L). For certain values of *d*, the shadow manifold M' is diffeomorphic, i.e., a 1:1 embedding of the original manifold (here for *d*=1, 12, 20 and 22, while for others it is not, due to crossings (here for *d*=2, 5, 15, 18 and 25) or collapse of the manifold (here for *d*=10). Note the similarity of the phase plots for every 10 steps of *d* (e.g. in panel D, H and L or in panel E vs. I and in panel C, G and K).

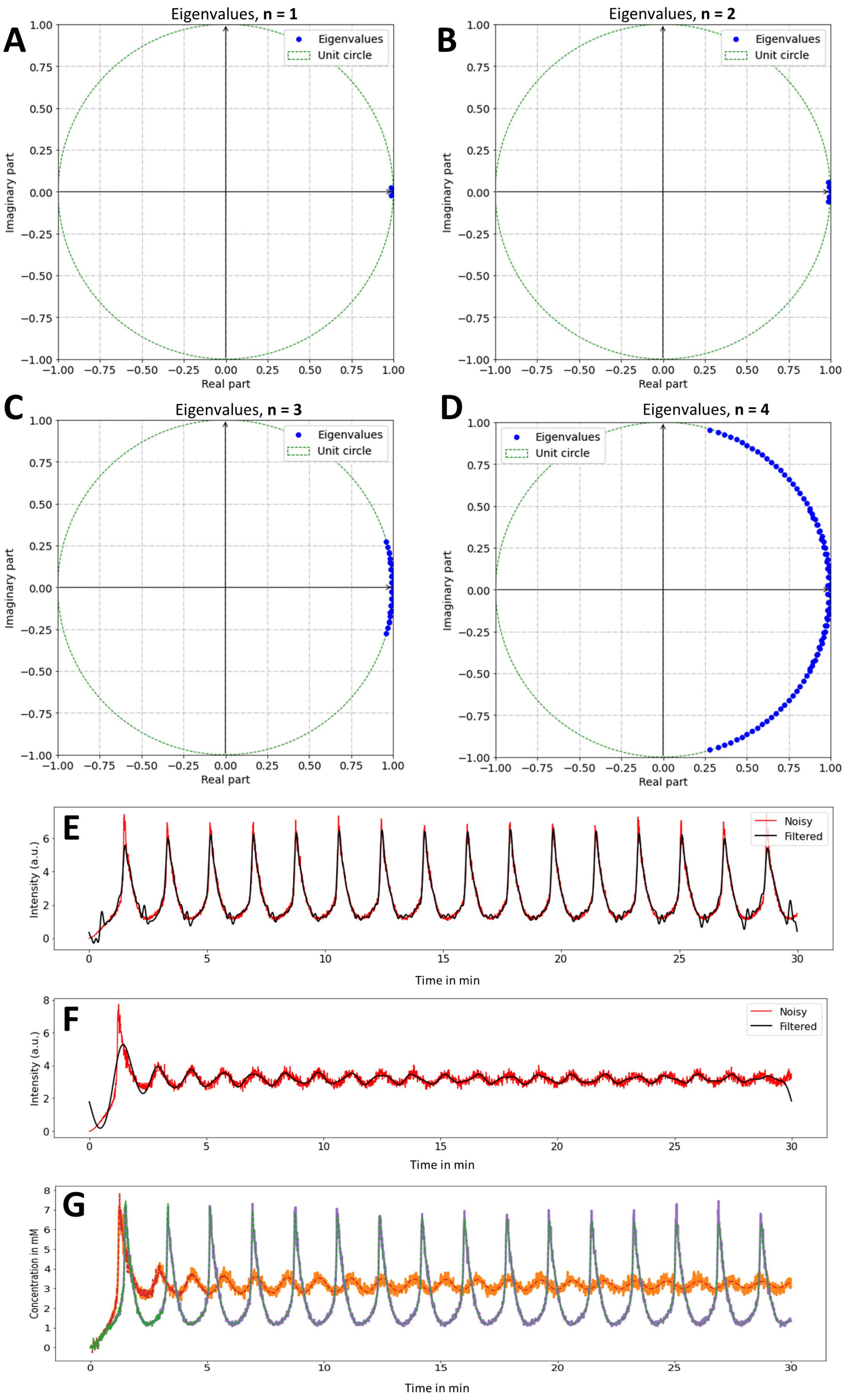


**Figure S2. Eigenvalues of HoDMD analysis and comparison with Fourier filtering.** The eigenvalue spectrum of the HoDMD analysis of the simulated time courses for varying Hill coefficients, *n*, is shown in A-D. E and F shows simulated time courses for F6P (E) and F16BP (F) with noise (red lines) and the resulting Fourier reconstruction (black lines). The reconstruction of the same data by HoDMD with rank truncation of r= 20 is shown in panel G for F6P (violet line, data; green line, reconstruction) and for F16BP (orange line, data; red line, reconstruction). See text for further details.

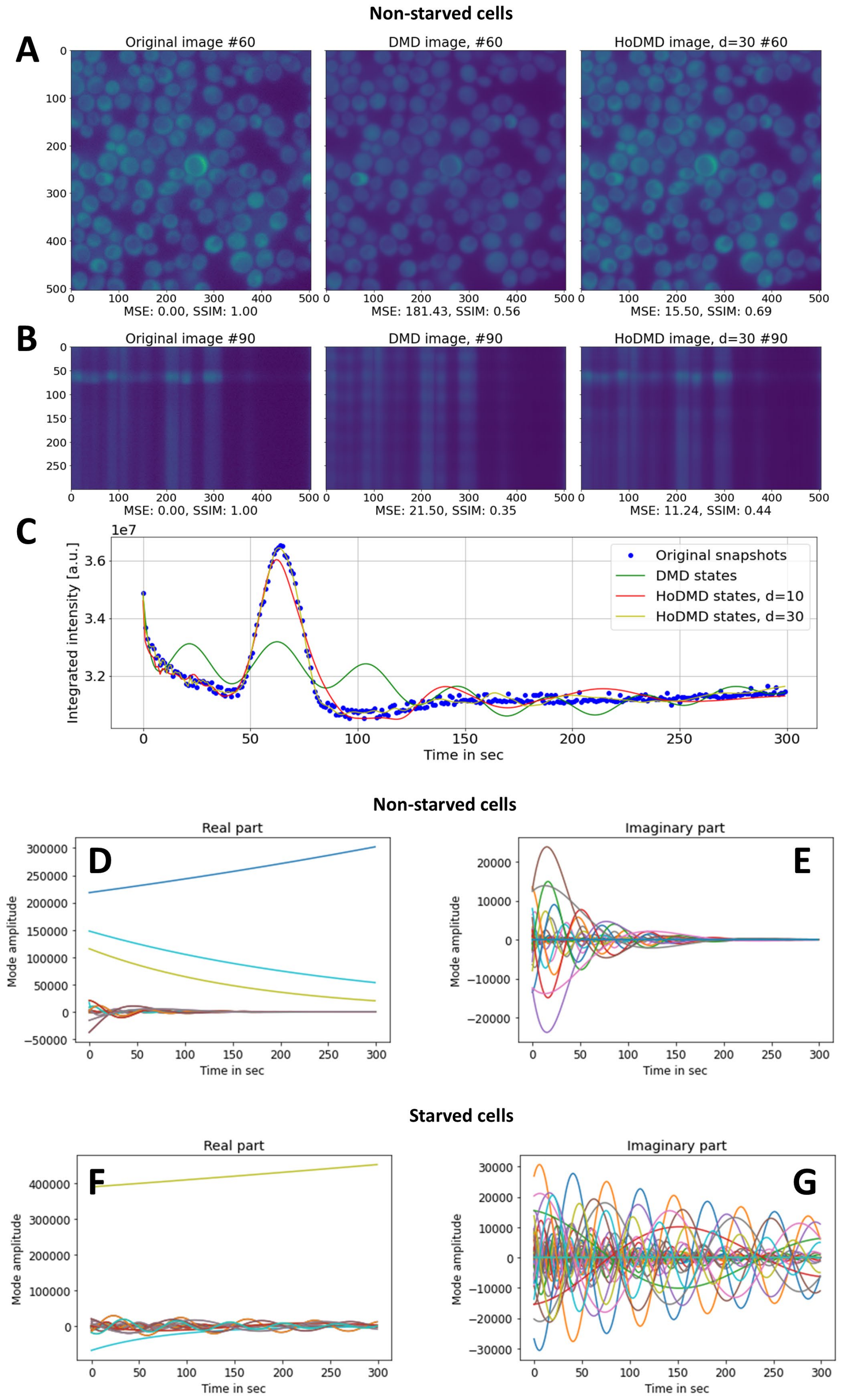


**Figure S3. HoDMD on image stacks of autofluorescence dynamics in non-starved yeast cells.** Cells were imaged on a wide field microscope every second, and the resulting image stacks were analyzed by standard DMD ('DMD') or by higher-order DMD ('HoDMD'). A, xy-view of one selected frame (frame 60); B, xz-profile along one selected line (line 90) for all three conditions. C, integrated intensity for the original snapshots (blue dots), the DMD reconstruction (green line) and the HoDMD reconstruction with either a delay of $d$ = 10 (red line) or $d$ = 30 (yellow line). Dynamics of the modes identified by HoDMD for this data, the non-starved cells, is shown in D (real part) and E (imaginary part). For comparison, the dynamics of the modes identified by HoDMD for starved cells (compare Fig. 6) is shown in F (real part) and G (imaginary part).

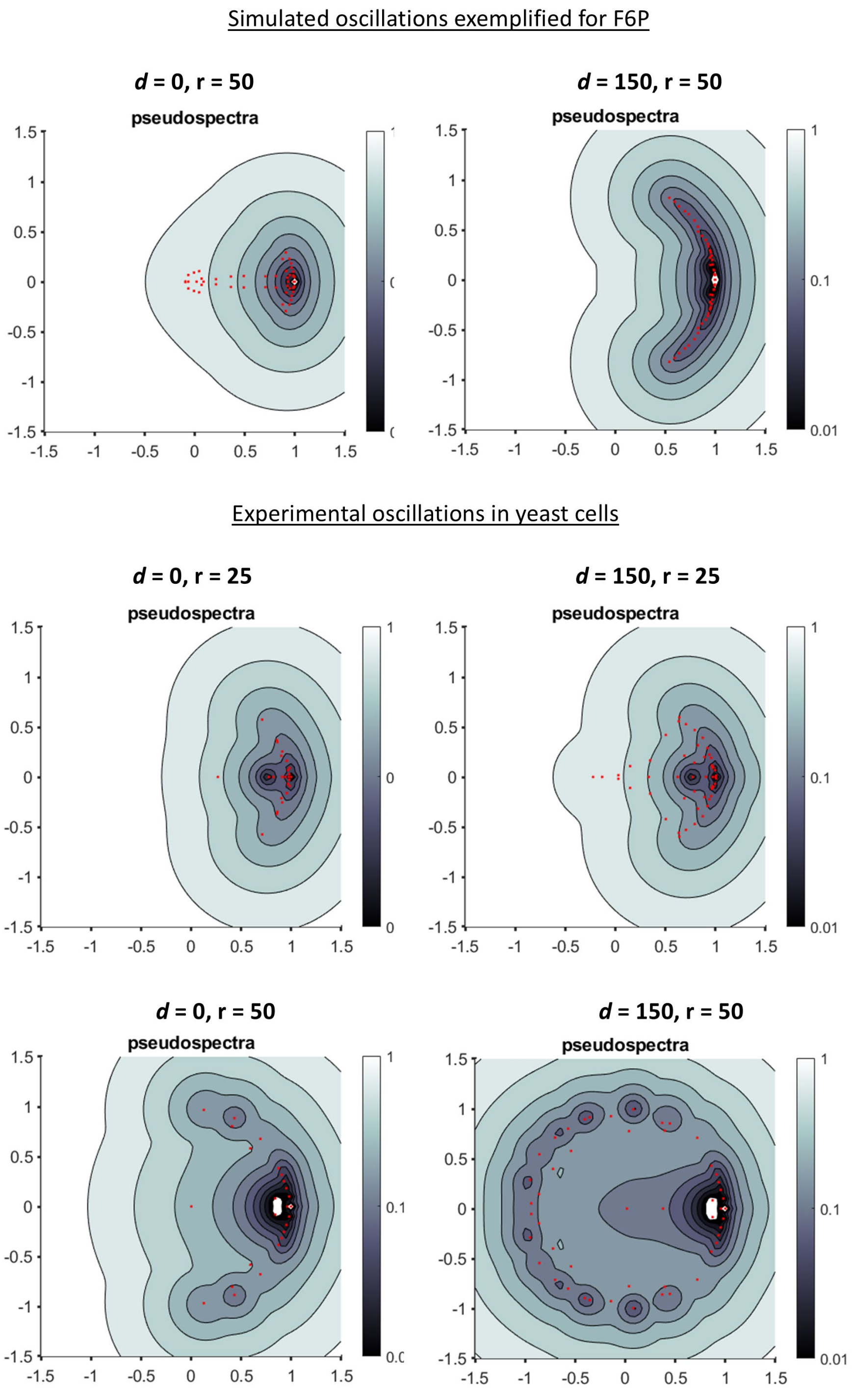


**Figure S4. Eigenvalue spectrum of synthetic and experimental oscillations by residual DMD.** Residual DMD was applied to cell-time matrices of the simulated data (upper panel) and of the experimental data (lower panel), either on the original data matrices without delay ($d$ = 0) or on the Hankel matrices ($d$ = 150) with rank truncation of r = 25 and r = 25 or 50 for the synthetic and experimental data, respectively. Red dots shows the identified eigenvalues on the unit circle, and grey shades are the associated residuals with dark grey indicating lower residuals and thereby higher confidence in the determined eigenvalues compared to light grey.

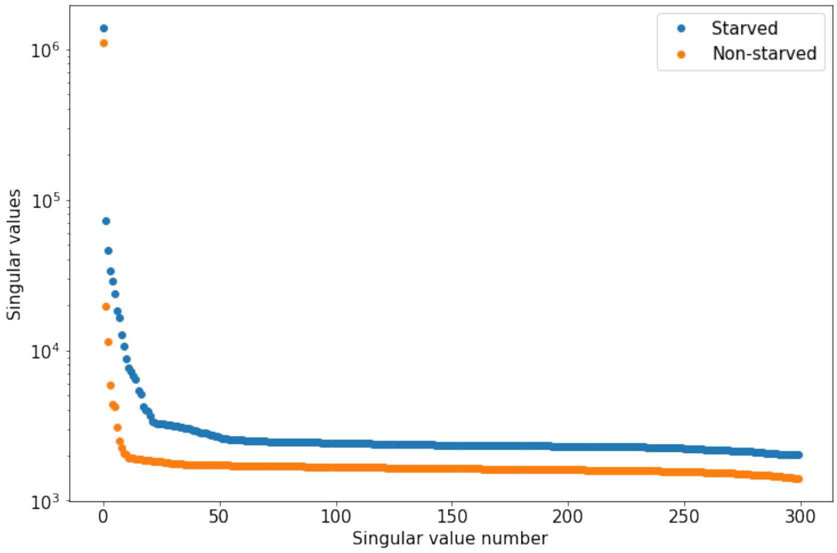


**Figure S5. Singular value spectrum of image data for starved and non-starved cells.** The singular values were calculated for the experimental data and plotted on a semi-logarithmic scale for starved (blue dots) and non-starved cells (orange dots).

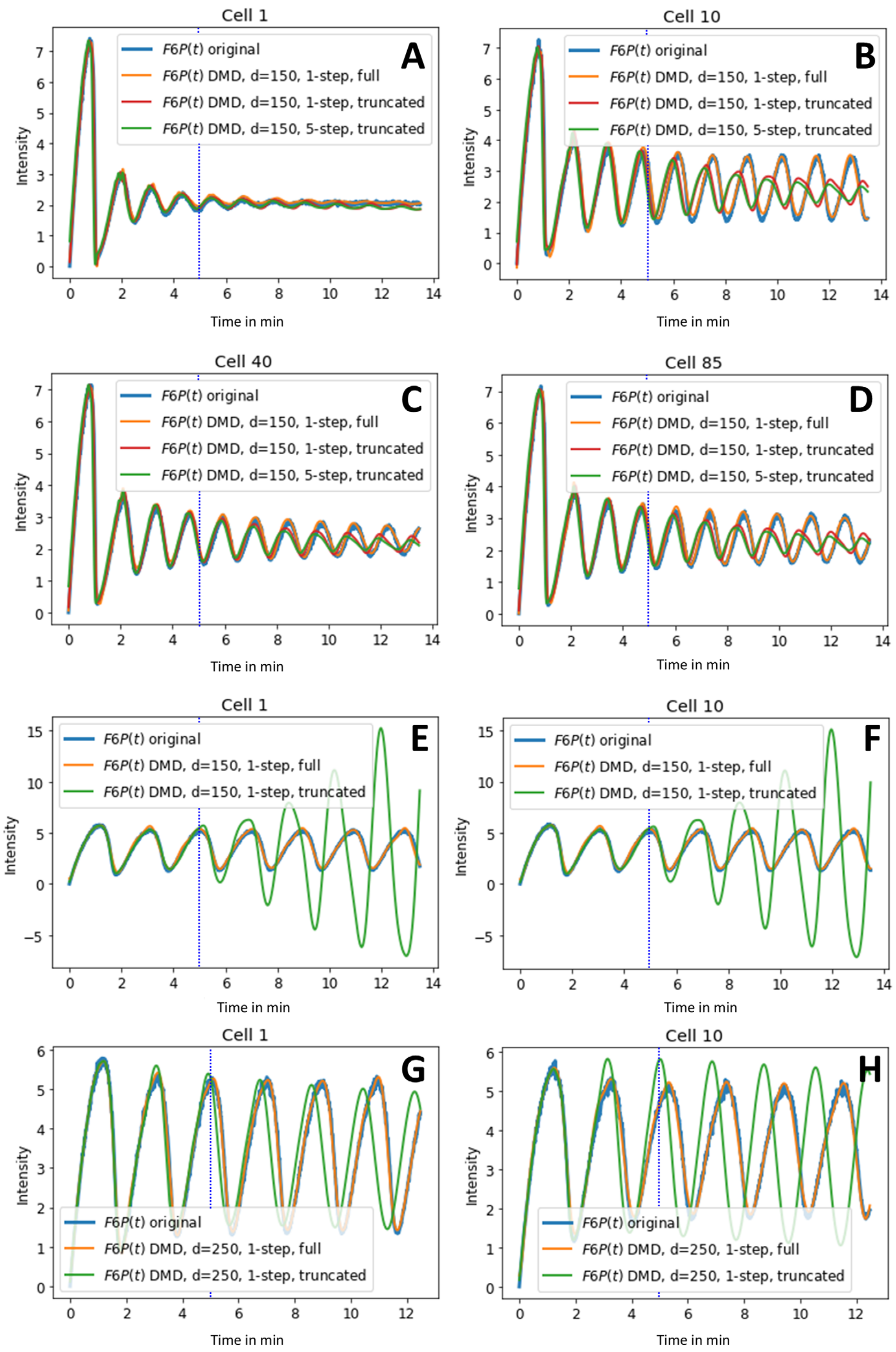


**Figure S6. Prediction capability of DMD with delay embedding on simulated time courses.** The ODE system of Eqs. 1 and 2 was simulated using 100 randomly generated values of glucose influx, v, as described in the main text, either with mean influx being 14.5 mM/min to get damped oscillations (A-D) or 9.5 mM/min to get sustained oscillation (E-H). Time courses for F6P are shown for selected cells as blue lines. DMD reconstructions with delay, *d* = 150 (A-F) or *d* = 250 (G-H) are shown. Color coding of the reconstructions is indicated in the legend of each panel. The vertical blue line shows the time point of truncation, left of which was used for training and right of which is the model prediction.

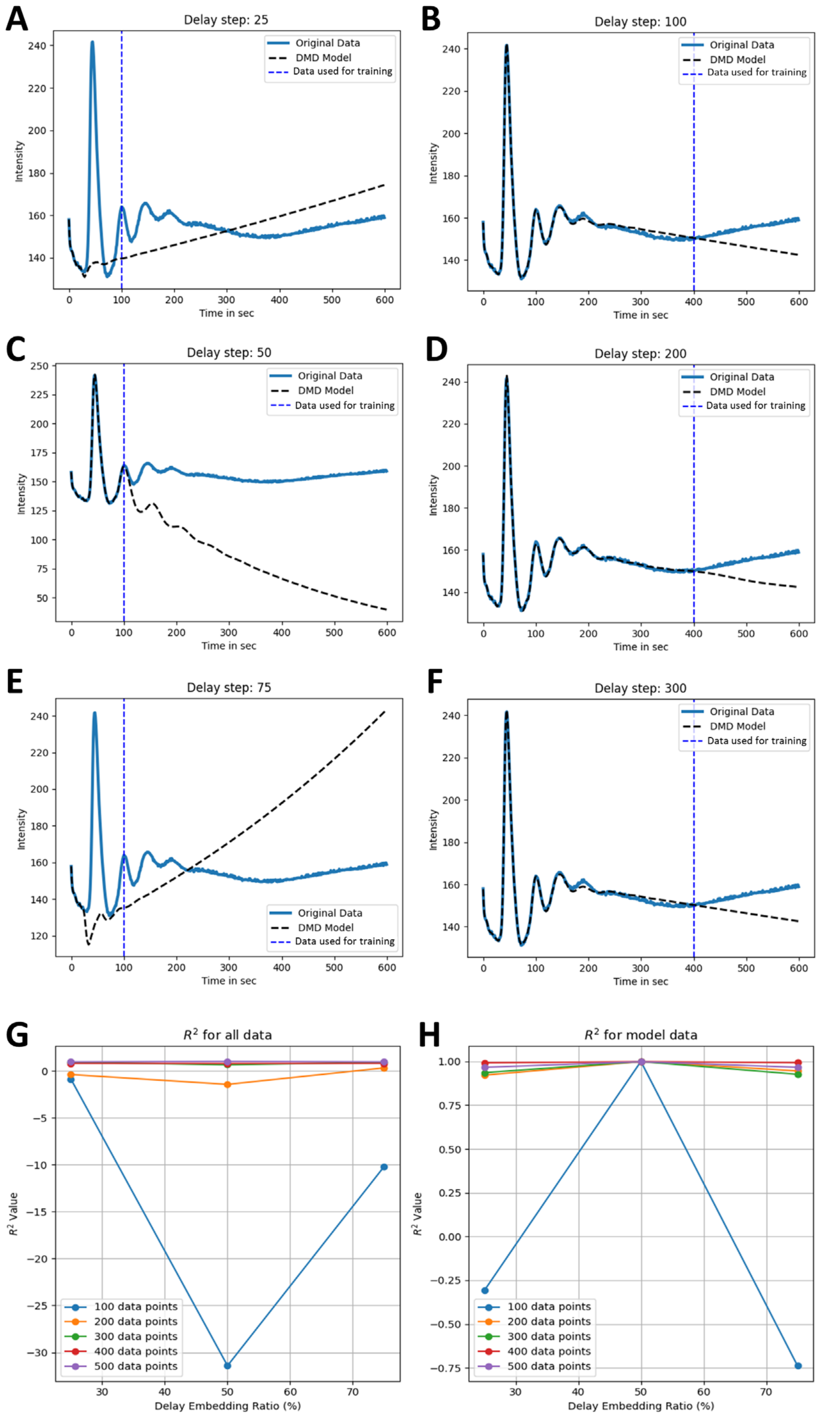


**Figure S7. Prediction of experimental time courses using DMD with time-delay embedding.** Experimental time courses of NADH fluctuations recorded in response to addition of 15 mM glucose were averaged for all cells in a field (blue lines in A-F, labeled 'Original data') and reconstructed using DMD with time-delay embedding (black dashed lines in A-F, labeled 'DMD model'). Training data started at t=0, and its length was varied with 100 time points in A, C and E, and 400 time points in B, D and F, as indicated by the dashed blue line in panel A-F (labeled 'Data used for training'). G and H shows the $R^2$-value calculated between either all data (G) or only training data (H) and the corresponding DMD reconstruction/prediction as function of delay, d for different lengths of the trainings data. See text for further explanations.

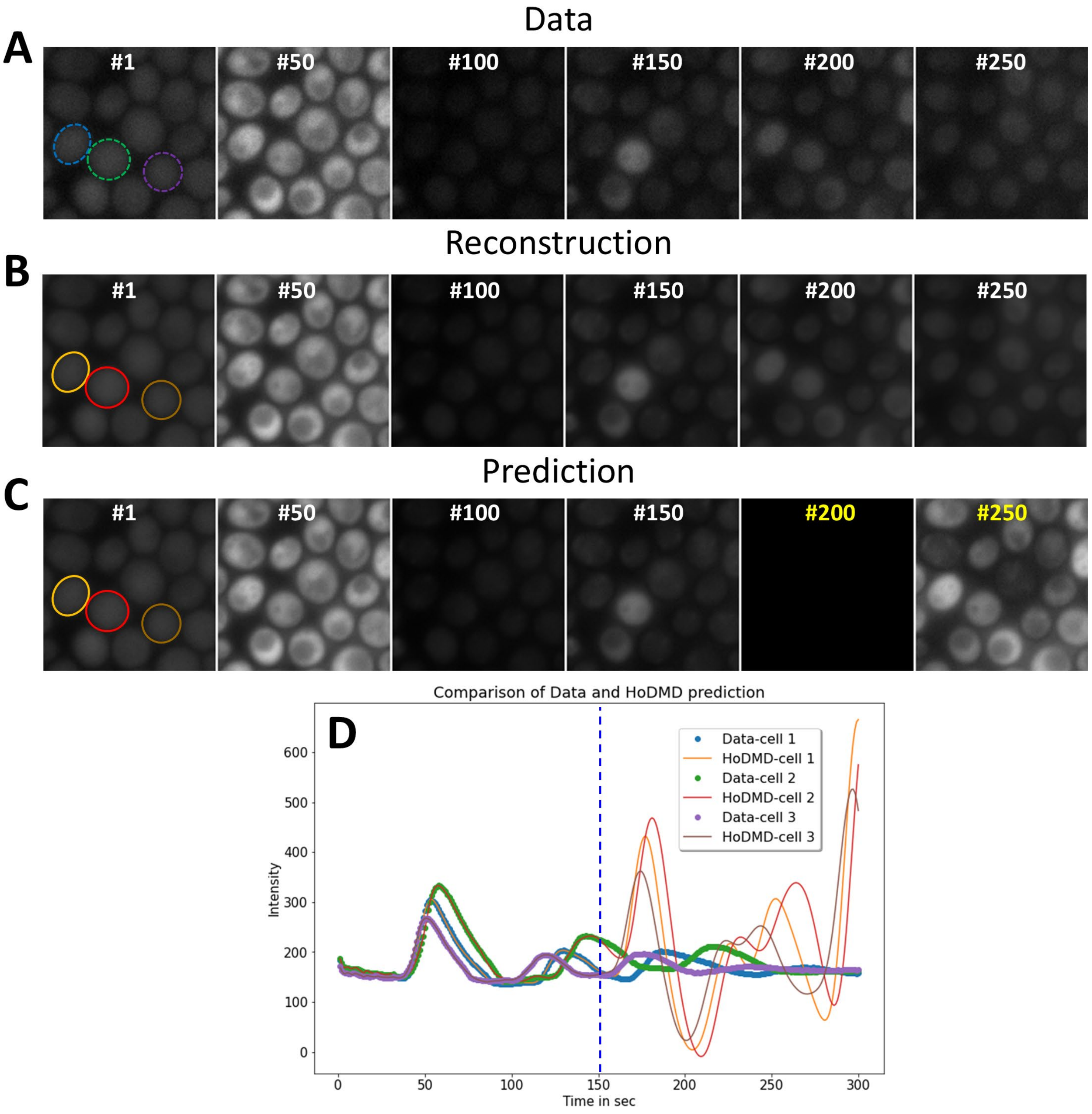


**Figure S8. Prediction of unseen images of yeast oscillations by HoDMD.** Image stacks of starved cells were analyzed by HoDMD, and a cropped region of the entire image data is shown (compare Fig. 6D-F). A-C, montage of every $50^{th}$ frame, starting from the first frame (#1) for the original data, A; the reconstruction, B; and the prediction; C. For B, the training data (white frame numbers) comprised the entire data set, while for panel C, training data are first 150 frames, while the last 150 frames are test data for predictions (yellow frame number). D, mean intensity profiles for the three cells outlined in A-C (cell 1, yellow; cell 2, red; cell 3, beige) with the blue dashed line indicating the border between training (left) and test data (right). Clearly, the prediction deviates rapidly from the data.

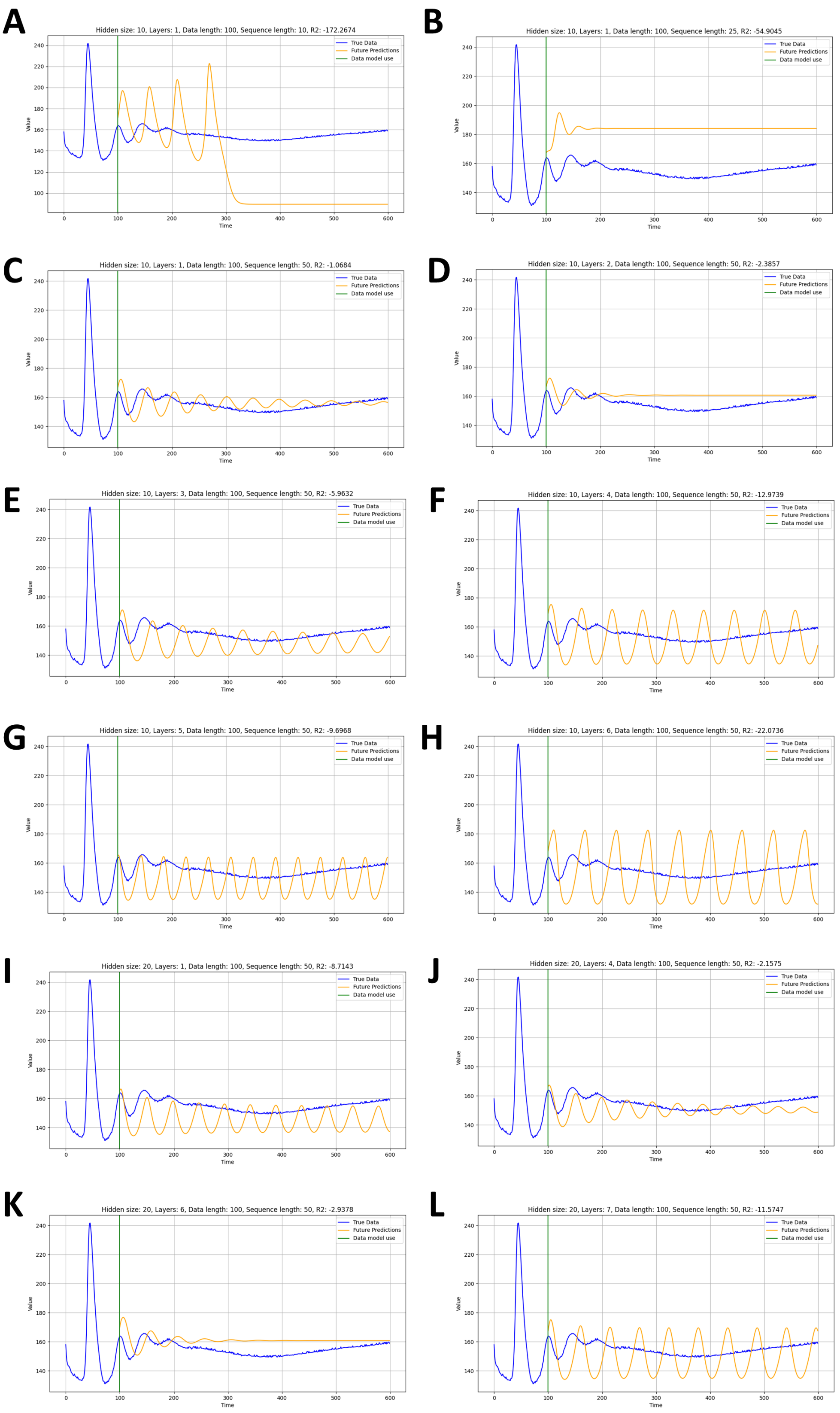


**Figure S9. Prediction of experimental time courses with LSTM network.** A LSTM was trained on the first 100 data points of the experimental autofluorescence time series (blue lines) to predict subsequent data points (yellow lines). Several network configurations were tested including the number of layers and nodes as well as the pecentage of training data included the sequence for training. Parameter combinations are given on top of each panel. The green line indicates the border between training data (0-100) and test data (101-600). See main text for further information.

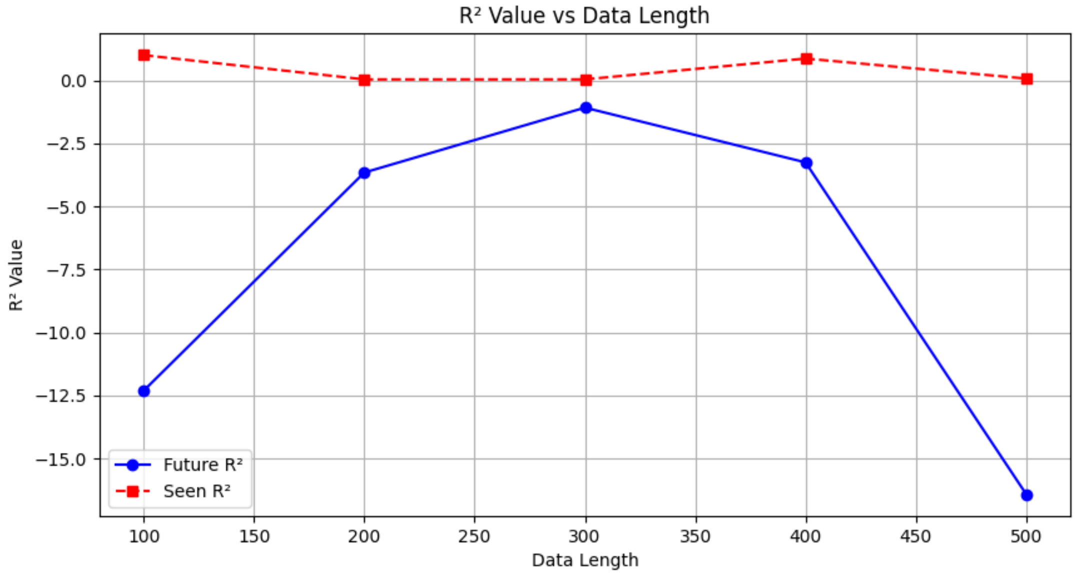


**Figure S10. LSTM performance as funcion of size of trainings data.** The LSTM network containing 4 hidden layers and 40 nodes with a sequence length of 50% of the data was trained for 200 epochs with the indicated length of training data, shown on the x-axis. The $R^2$-value was calculated separately for the performance on the training data ('Seen $R^2$', red) and the test data (Future $R^2$', blue).
